# Identification of different intrinsic sequence patterns between HIV-1 DNA and RNA across subtypes using the *k*-mer-based approach

**DOI:** 10.64898/2026.02.25.707904

**Authors:** Janusz Wiśniewski, Karol Serwin, Miłosz Parczewski, Anna Kula-Păcurar, Pavel Skums, Alexander Kirpich, Sergiy Yakovlev, Heng-Chang Chen

## Abstract

Advanced analytical tools that enable mining of the masked features hidden in intricate datasets and strengthening the biological interpretation of multigenomic outputs hold paramount importance. At present, HIV-1 subtyping remains a challenging task in a great part due to analytical tool discordance. To tackle this issue, in this study, we present an updated version of a *k*-mer-based approach, PORT-EK-v2, a streamlined bioinformatic pipeline, allowing for a comparison of multiple genomic datasets and identification of over-represented genomic regions, *k*-mers, related to specific origins of datasets. Using PORT-EK-v2, we exemplified that intrinsic sequence patterns between HIV-1 DNA and RNA are distinct across group M HIV-1 subtypes. Furthermore, we showcased that “isolate *k*-mer count”, a predictive variable computed in this work, could serve as a default choice in classifying the HIV-1 DNA versus RNA sequences across subtypes. Lastly, results based on network-based analyses and Markov chain Monte Carlo modeling unveiled a clear discontinuation of a random walk throughout the network properties corresponding to each tested group of HIV-1 subtypes, confirming the specificity of enriched *k*-mer retrieved by PORT-EK-v2 and the genomic diversity across group M HIV-1 subtypes. Source code for PORT-EK-v2 is at https://github.com/Quantitative-Virology-Research-Group/PORT-EK-version-2 and is freely available.

## Introduction

The current human immunodeficiency type 1 (HIV-1) pandemic is phylogenetically classified into M, N, O, and P four groups^1–3^ based on RNA sequences. Among them, group M is the most prevalent worldwide and is responsible for pandemics in humans. This high genomic diversity is one of the main characteristics of the HIV-1 genomes due to its high mutation rate, a lack of proofreading activity of reverse transcriptase^4,5^, viral recombination-prone nature^6–8^, a lack of proofreading of its replicated RNA genomes^9^, and alternative splicing^10,11,12^. Nevertheless, HIV-1 RNA is crucial for tracking the transmission dynamics of primary HIV-1 infections, monitoring the circulation and evolution of HIV-1 subtypes and variants, and detecting emerging drug-resistance mutations in HIV-1.

In cases where the viral load is too low for successful HIV-1 RNA genotyping, HIV-1 DNA may serve as an alternative to identify drug resistance mutations within patients subjected to antiretroviral therapy (ART), as evidenced by several studies^13-17^. Although the feasibility of the proviral DNA-based HIV-1 resistance testing remains debated, this strategy often captures apolipoprotein B mRNA editing enzyme, catalytic polypeptide-like (APOBEC)-mediated G-to-A hypermutation and genetically defective and replication-incompetent viruses^18-20^. Indeed, more than 90% of proviral sequences are genetically defective^21^, particularly falling in the region of the HIV *env* gene^22,23^, further exacerbating the heterogeneous nature of proviral genomes in reservoir cells. Although diversities between HIV-1 DNA and RNA have been previously characterized, appropriate HIV-1 subtyping remains an unsolved issue. Given that the composition of a genomic sequence can be viewed at different hierarchical orders, ranging from nucleotides, sequence motifs, individual genes, to a cluster of genes involved in the same regulatory network, in this work, we investigate the HIV-1 DNA and RNA genomic sequence diversity at the level of enriched *k*-mers, short genomic sequences [i.e., 13 base pairs (bp), 15 bp, and 17 bp], and hypothesize that the difference in the intrinsic sequence pattern between HIV-1 DNA and RNA exists and may influence the classification of HIV-1 genomic diversity. More comprehensive insights into the difference between HIV-1 DNA and RNA genomic sequences may enhance the accuracy and the granularity of genome surveillance of HIV-1^24,25^.

Traditional methods for HIV-1 taxonomies are mainly based on gene-based phylogenetic analysis embedded with alignment-based methods. Individual HIV-1 gene sequences (often from the *env* gene^26^ and *gag* region^27^) are aligned with curated subtype reference sequences, followed by a comparison of homologous nucleotide patterns or motifs to create evolutionary trees^28,29^. Such methods can be computationally expensive and time-consuming, given that similarity statistics over a sliding window are computed^30,31^. In addition, presently available reference genomes may not be appropriate for all subspecies or subtypes within certain species, resulting in inaccurate alignment results.

To conquer this limitation, various alignment-free classification methods have been proposed. COMET (COntext-based Modeling for Expeditious Typing)^32^ was developed based on adaptive coding and partial string matching^33^. CASTOR^34^ was developed based on restriction fragment length polymorphism (RFLP) and has been applied to the subtyping of human papillomavirus (HPV), hepatitis B virus (HBV), and HIV-1. The “Natural Vector representation” approach that classifies single-segmented^35^ and multi-segmented^36^ viral genomes, as well as viral proteomes^37^. Phymm, an interpolated Markov-based model, has been applied to the phylogenetic classification of metagenomic samples^38^.

In addition to the mentioned methods, the use of *k*-mers is also employed for phylogenetic applications. Various *k*-mer-based alignment-free methods, including K^AMERIS39^, a multi-label learning algorithm^40^, KINN^41^, KEVOLVE^42^, and a *k*-mer-based eXtreme Gradient Boosting model^43^, have been proposed for HIV-1 subtyping and phylogenetic analysis^41,44,45^, and HIV-1-resistent drug detection^46^. Although the feasibility of *k*-mer-based methods may mitigate the difficulty in the classification of HIV-1 subtypes, limitations exist. For example, while K^AMERIS39^ achieves high overall accuracy, recall across minority classes seems to be insufficient. Although KEVOLVE^42^ achieves good classification performance, it tends to mistakenly classify recombinant subtypes as pure subtypes^43^. The *k*-mer and position-based vectorization method, in conjunction with multi-class k-nearest neighbours (KNN) algorithm, was developed using only a single gene, which may risk failing to capture the complexity of the entire HIV-1 genome^40^.

In this work, we showcased the enhanced effectiveness and accuracy of the streamlined pipeline of the Pathogen Origin Recognition Tool using Enriched *K*-mers (PORT-EK)^47^ version 2, namely PORT-EK-v2. PORT-EK-v2 is a *k*-mer-based approach, which facilitates the comparison of various multigenomic datasets of HIV-1 subtypes and the detection of over-represented genomic regions (i.e., *k*-mers) linked to specific sequences. Our contributions to PORT-EK-v2 are two fold. First, PORT-EK-v2 computed the relative abundance of enriched *k*-mers (i.e., enriched *k*-mer frequency) between two compared multigenomic datasets, aiming at identifying “quantitative” differences between two species. This feature in PORT-EK-v2 enables unveiling more masked patterns previously hidden in intricate datasets, subsequently gaining a more profound interpretation of results. Second, PORT-EK-v2 automatically computes the optimal length of *k*-mers fitting to the input dataset and generates *k*-mer frequency enrichment statistics in a readable table format (see **Supporting information**). These features allow PORT-EK-v2 to outperform other similar methods. In this work, we applied PORT-EK-v2 to pinpoint *k*-mers abundance in HIV-1 DNA or RNA sequences and showcased distinct sequence patterns between HIV-1 DNA and RNA across group M HIV-1 subtypes.

## Results

### An overview of PORT-EK-v2

The critical steps of PORT-EK-v2 include (i) *k*-mers matrix preparation, (ii) *k*-mers filtering and selection of enriched *k-*mers, and (iii) mapping of enriched *k*-mers to a reference genome. The predictive accuracy using *k*-mer-related features was tested independently from the PORT-EK-v2 pipeline. Compared with PORT-EK^47^, technical advances in PORT-EK-v2 include (i) the establishment of a streamlined computing pipeline, allowing for reducing computing cost (computational resources requirement) (**Table 1**) and offering minimal computing time (**Table 1**), and leveraging its effectiveness, and (ii) the improvement of mapping algorithms, thereby providing a superior performance of retrieving enriched *k*-mers. Detailed rationale and background of PORT-EK-v2 and its optimized parameterization in comparison to PORT-EK^47^ are provided in **Supporting information**. The systematic comparison of key parameters between PORT-EK^47^ and PORT-EK-v2 is summarized in **Table S1**. In addition, using input HIV-1 RNA sequences and two subsets of randomly bootstrapped HIV-1 RNA sequences of different sizes, we benchmarked PORT-EK-v2 against PORT-EK^47^, revealing a remarkable reduction in computing speed and peak memory footprint of the enrichment of *k*-mer frequencies (**Table 1**). In addition, the *k*-mer counting speed of PORT-EK-v2 was also evaluated to be comparable to that achieved by Jellyfish^48^, Kmer-db^49,50^, and KMC 3^51^ (**Table 2**). It is important to stress that PORT-EK^47^ and PORT-EK-v2 are designed and optimized for a large number of small genomes (mainly for viral infectious diseases), as it calculates aggregate statistics concurrently with *k*-mer counting, which may lead to a slow progression for large genomes.

**Table 1.**
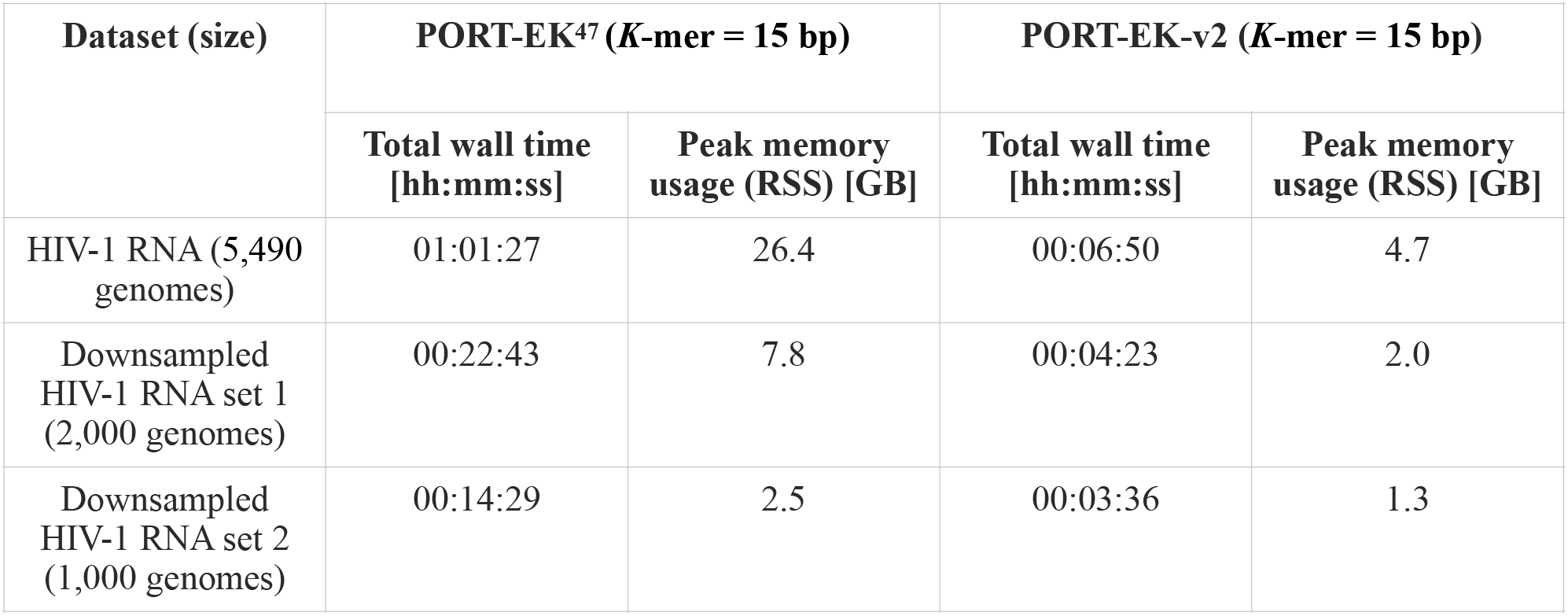
Benchmarking results of PORT-EK-v2 versus PORT-EK.

| **Dataset (size)** | **PORT-EK<sup>47</sup> (*K*-mer = 15 bp)** |  | **PORT-EK-v2 (*K*-mer = 15 bp)** |  |
| --- | --- | --- | --- | --- |
|  | **Total wall time**<br>**[hh:mm:ss]** | **Peak memory**<br>**usage (RSS) [GB]** | **Total wall time**<br>**[hh:mm:ss]** | **Peak memory**<br>**usage (RSS) [GB]** |
| HIV-1 RNA (5,490<br>genomes) | 01:01:27 | 26.4 | 00:06:50 | 4.7 |
| Downsampled<br>HIV-1 RNA set 1<br>(2,000 genomes) | 00:22:43 | 7.8 | 00:04:23 | 2.0 |
| Downsampled<br>HIV-1 RNA set 2<br>(1,000 genomes) | 00:14:29 | 2.5 | 00:03:36 | 1.3 |

**Table 2.**
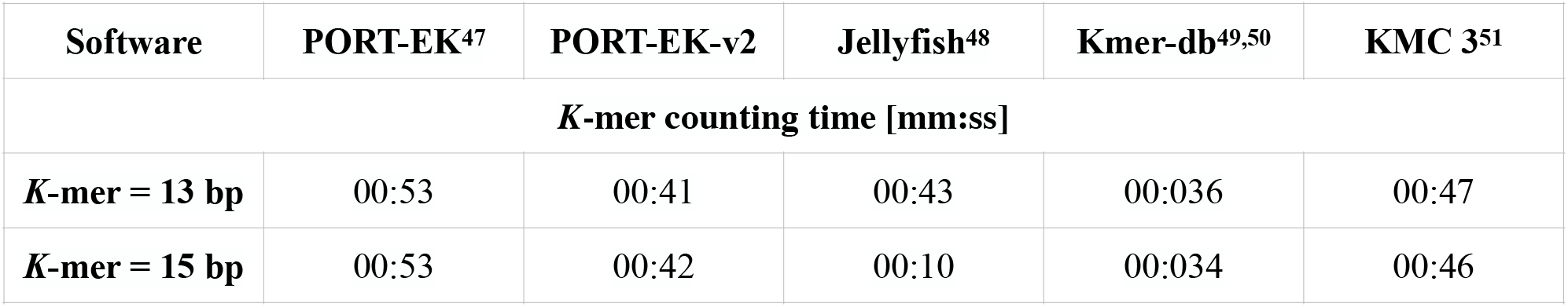
Benchmarking results of *k*-mer counting speed using PORT-EK-v2, Jellyfish, Kmer-db, and KMC 3 based on HIV-1 RNA sequences (5,490 genomes).

| Software | PORT-EK<sup>47</sup> | PORT-EK-v2 | Jellyfish<sup>48</sup> | Kmer-db<sup>49,50</sup> | KMC 3<sup>51</sup> |
| --- | --- | --- | --- | --- | --- |
| **$K$-mer counting time [mm:ss]** |  |  |  |  |  |
| **$K$-mer = 13 bp** | 00:53 | 00:41 | 00:43 | 00:036 | 00:47 |
| **$K$-mer = 15 bp** | 00:53 | 00:42 | 00:10 | 00:034 | 00:46 |

### Distinct sequence patterns unveiled by enriched *k*-mers between HIV-1 DNA and RNA across subtypes

A total of 10,015 HIV-1 DNA and 5,494 RNA sequences were downloaded from the Los Alamos National Laboratory HIV Databases (https://www.hiv.lanl.gov/). After the examination of the quality of genomic sequences, the subsets of 10,013 and 5,490 HIV-1 DNA and RNA sequences were subject to PORT-EK-v2 and downstream analyses. Analyzed sequences and corresponding exact IDs are archived at GitHub (see **Methods**). To minimize a bias resulting from the imbalance of the sample size across each subtype, we combined genomes from rare subtypes for which a small number of sequences were retrieved as one group named “rare subtypes” for the following analysis. Eventually, five groups of the group M HIV-1 subtypes— the subtypes A, B, C, D, and the rare subtypes, including subtypes F1, F2, G, H, J, K, and L—were computed in this work. To identify enriched *k*-mers within a specific HIV-1 subtype, the “*k*-mer average count” (refer to **Supporting information**) has to be substantially greater in one group than in others based on pairwise comparisons (detailed in **Supporting information**).

Using PORT-EK-v2, we first observed distinct distributions between enriched DNA and RNA *k*-mers with a length of 13 bp [hereinafter, total enriched DNA *k*-mers (13 bp), **Fig. 1a**; total enriched RNA *k*-mers (13 bp), **Fig. 1c**] and 15 bp [hereinafter, total enriched DNA *k*-mers (15 bp), **Fig. 1b**; total enriched RNA *k*-mers (15 bp), **Fig. 1d**]. Based on enriched DNA *k*-mers, a bipolar distribution with subtypes B and C was observed (**Fig. 1a** and **1b**). A linear distribution of enriched DNA *k*-mers, aligning with subtypes C, A, the rare subtypes, D, and B, reflects a linear similarity of DNA sequence in terms of enriched *k*-mer frequencies across isolates in different subtypes. DNA sequences retrieved from subtypes A, D, and the rare subtypes shared a relatively high sequence similarity compared with sequences retrieved from subtypes B and C (**Fig. 1a** and **1b**). Of note, a small fraction of enriched DNA *k*-mers in subtype B was observed. In contrast, a tripolar and an equivalent radial distribution of enriched RNA *k*-mers inferred that quantitatively, the RNA sequence diversity of isolates across subtypes A, B, and C is indistinguishable (**Fig. 1c** and **1d**). The relative position of subtype A genomes is, however, markedly different from subtypes B and C (**Fig. 1c** and **1d**). The similarity of RNA sequences was shared between subtypes B and D, as well as subtype D and the rare subtypes. Similar distribution patterns also appeared in enriched DNA *k*-mers (17 bp) (**Supplementary Fig. S1a**) and enriched RNA *k*-mers (17 bp) (**Supplementary Fig. S1b**), suggesting the substantial differences in sequence patterns between HIV-1 DNA and RNA.

**Fig. 1.**
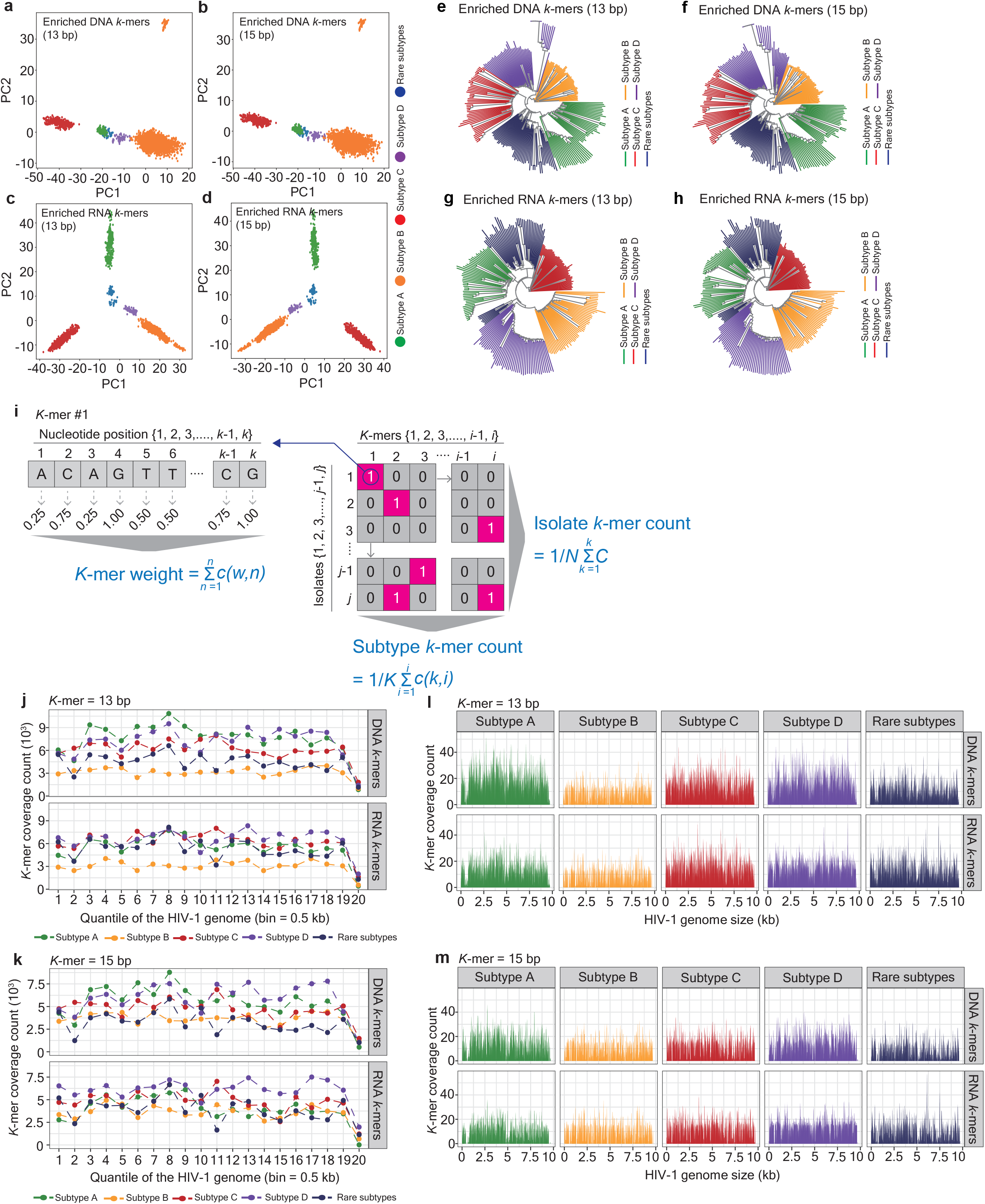
Distinct enriched DNA and RNA *k*-mers across different groups of HIV-1 subtypes. (**a-d**) Two dimensional principal component analysis (PCA) revealing the discrepancy of enriched DNA (**a**, *k*-mer = 13 bp; **b**, *k*-mer = 15 bp) and RNA *k*-mers (**c**, *k*-mer = 13 bp; **d**, *k*-mer = 15 bp). Dots marked green, orange, red, purple, and dark blue represent HIV-1 subtype A, B, C, D, and rare subtypes, respectively. (**e-h**) Phylogenetic trees representing the evolutionary distance across isolates assigned to different groups of HIV-1 subtypes. Trees were constructed by “*k*-mer average count” computed from enriched DNA (**e**, *k*-mer = 13 bp; **f**, *k*-mer = 15 bp) and RNA (**g**, *k*-mer = 13 bp; **h**, *k*-mer = 15 bp) *k*-mers across different groups of HIV-1 subtypes. 250 enriched *k*-mers were bootstrapped from each pool of enriched *k*-mers to illustrate a tree plot. The color code annotation denotes the group of HIV-1 subtypes: green, orange, red, purple, and dark blue represent HIV-1 subtype A, B, C, D, and rare subtypes, respectively. (**i**) Schematic representation of the definitions of enriched *k*-mer-related features, including “*k*-mer weight”, “subtype *k*-mer count”, and “isolate *k*-mer count”. Corresponding mathematical equations were provided along with respective features. The definition of each parameter can be found in **Methods** and **Supporting information**. (**j, k**) Quantile plots representing the coverage of enriched *k*-mers in a length of 13 (**j**) or 15 bp (**k**) throughout the complete HIV-1 genome at the quantile of 0.5 kb. Facets on the y-axis separate enriched *k*-mers computed by the DNA or RNA sequence. Dots and dashed lines marked green, orange, red, purple, and dark blue represent HIV-1 subtype A, B, C, D, and rare subtypes, respectively. (**l, m**) Line plots representing the coverage of enriched *k*-mers in a length of 13 (**l**) or 15 bp (**m**) throughout the complete HIV-1 genome. Facets at the x-axis separate different groups of HIV-1 subtypes; facets at the y-axis separate enriched DNA and RNA *k*-mers. Lines marked green, orange, red, purple, and dark blue represent HIV-1 subtype A, B, C, D, and rare subtypes, respectively.

A clear separation across different groups of HIV-1 subtypes was also confirmed by phylogenetic trees, in which the distance is computed based on enriched *k*-mer count (**Fig. 1e-1h**). Intriguingly, a small subset appearing in subtypes A and D inferred the presence of a higher diversity in enriched DNA *k*-mer count in these two groups compared with others (**Fig. 1e** and **1f**), whereas an independent subset was detected in the rare subtypes when enriched RNA *k*-mers were processed (**Fig. 1g** and **1h**). No significant difference in the proportion of enriched *k*-mers that are exclusively present in corresponding groups can be detected (**Supplementary Fig. S1c**).

We further computed three additional features of enriched *k*-mers: (1) “*k*-mer weight”—representing the composition of each enriched *k*-mer at a nucleotide level, (2) “subtype *k*-mer count”—the sum of individual *k*-mers normalized by different groups of HIV-1 subtypes, and (3) “isolate *k*-mer count”—the appearance of individual *k*-mers at a single isolate level (**Fig. 1i**). The definition of each feature is detailed in **Methods**. We observed a modest and significant enrichment of enriched DNA and RNA *k*-mers in subtype C compared with *k*-mers enriched in other groups of HIV-1 subtypes while comparing “*k*-mer weight” across each group (**Supplementary Fig. S1d**). No difference in “*k*-mer weight” between *k*-mers in a length of 13 bp and 15 bp, except for a minor enrichment in enriched DNA *k*-mers (13 bp) in subtypes B (enriched DNA *k*-mers: 7.564, enriched RNA *k*-mers: 7.501) and C (enriched DNA *k*-mers: 7.463, enriched RNA *k*-mers: 7.416) compared with enriched RNA *k*-mers (**Supplementary Fig. S1e**). Although a bias was observed when different ordinal encoding was assigned to every nucleotide in an enriched *k*-mers (**Supplementary Fig. S2a-S2c**), the overall patterns of “*k*-mer weight” across different groups of HIV-1 subtypes were reproducible (**Supplementary Fig. S2d-S2f**).

A more abundant “subtype *k*-mer count” appeared in subtype D, whereas relatively few counts were in subtype B, irrespective of enriched DNA or RNA *k*-mers and their lengths (**Supplementary Fig. S1f**). A diverse and varied “subtype *k*-mer count” between enriched DNA and RNA *k*-mers across different groups of HIV-1 subtypes was also observed (**Supplementary Fig. S1g**).

While computing “isolate *k*-mer count”, a superior enrichment of enriched DNA *k*-mers appeared in the group of rare subtypes, followed by subtype D and A (**Supplementary Fig. S1h**), whereas the majority of enriched RNA *k*-mers accumulated in the HIV-1 subtype D, followed by the rare subtypes and a relatively minor number of *k*-mers were enriched in isolates across the subtypes A, B, and C (**Supplementary Fig. S1h**). A higher “isolate *k*-mer count” in enriched DNA *k*-mers than in enriched RNA *k*-mers was observed in subtypes A, C, and the rare subtypes (**Supplementary Fig. S1i**), whereas an opposite pattern was detected in subtypes B and D (**Supplementary Fig. S1i**).

To visualize the coverage of enriched DNA and RNA *k*-mers, we summarized the HIV-1 genome by 20 different quantile values (**Fig. 1j**, *k*-mer = 13 bp; **Fig. 1k**, *k*-mer = 15 bp) and observed that a higher enriched *k*-mer coverage appears in subtypes A and D, followed by subtype C and the rare subtypes, whereas the coverage of enriched RNA *k*-mers across subtypes A, C, D, and the rare subtypes was comparable (**Fig. 1j** and **1k**). Identical patterns were observed in the circumstances that the HIV-1 genome was summarized by 10 (**Supplementary Fig. S1j**, *k*-mer = 13 bp; **Supplementary Fig. S1k**, *k*-mer = 15 bp) and 7 (**Supplementary Fig. S1l**, *k*-mer = 13 bp; **Supplementary Fig. S1m**, *k*-mer = 15 bp) different quantile values. The overall coverage was illustrated in **Fig. 1l** (*k*-mer = 13 bp) and **1m** (*k*-mer = 15 bp). The coverage of enriched RNA *k*-mers of length 15 bp in subtype D was slightly higher than in other groups of HIV-1 subtypes. In all scenarios, a low *k*-mer coverage was shown in subtype B. Overall, these observations suggested that the sequence pattern varies between HIV-1 DNA and RNA sequences and differs across HIV-1 subtypes, as unveiled by enriched *k*-mers.

### Distinct genomic compositions between HIV-1 DNA and RNA were revealed at the sequence level rather than at a genic level

Next, we retrieved enriched *k*-mers that are exclusively present in either DNA (hereinafter, unique enriched DNA *k*-mers) or RNA (hereinafter, unique enriched RNA *k*-mers) enriched *k*-mers, separated by distinct groups of HIV-1 subtypes (**Supplementary Fig. S3a**). We computed that a range between 15% to 40% of unique enriched DNA *k*-mers and 15% to 39% of unique enriched RNA *k*-mers were present in corresponding groups of HIV-1 subtypes (**Supplementary Fig. S3a**). Distinct magnitudes of *k*-mer enrichment (measured by “*k*-mer RMSE”, see **Supporting information**) verified a strong correlation between unique enriched *k*-mers and distinct genomic composition across different groups of HIV-1 subtypes (**Supplementary Fig. S3b**).

As we computed “*k*-mer weight”, only minor differences across different groups of HIV-1 subtypes were observed (**Supplementary Fig. S3c**). Notably, unlike the pattern shown in total enriched DNA *k*-mers (**Supplementary Fig. S1f**), the increased abundance of unique enriched DNA *k*-mers accumulated in subtype B, highlighting the possibility that subgenomic locations throughout the DNA sequence (unique versus common *k*-mers) are present, as “subtype *k*-mer count” was examined (**Supplementary Fig. S3d**). In contrast, a similar pattern between total and unique enriched RNA *k*-mers was observed (**Supplementary Fig. S1f** and **Fig. S3d**).

To target unique enriched *k*-mers with significant enrichment, we retrieved those with the fold change in RMSE greater than 0.5 (arbitrary units) (**Fig. 2a**, *k*-mer = 13 bp; **Fig. 2c**, *k*-mer = 15 bp) and 1 (**Fig. 2b**, *k*-mer = 13 bp; **Fig. 2d**, *k*-mer = 15 bp) in a logarithmic scale (log^2^). The majority of unique enriched DNA *k*-mers appeared in subtype A, whereas abundant unique enriched RNA *k*-mers accumulated in subtype D (**Fig. 2e**). This observation exemplified that the inconsistency between HIV-1 DNA and RNA within the same group of HIV-1 subtypes exists. Similar patterns were observed in both cutoff thresholds. We also depicted the landscapes of unique enriched DNA and RNA *k*-mers throughout the HIV-1 genome, verifying the phenotype that distinct loci overlapping unique enriched DNA and RNA *k*-mers across different groups of HIV-1 subtypes were present (**Supplementary Fig. S3e**, *k*-mer = 13 bp; **Supplementary Fig. S3f**, *k*-mer = 15 bp). A superior frequency of unique enriched *k*-mers overlapped the HIV-1 *pol* gene, followed by the *env* and *gag* genes (**Fig. 2f**, *k*-mer = 13 bp; **Fig. 2g**, *k*-mer = 15 bp). Such a bias was, however, indistinguishable either between DNA and RNA sequences or across different groups of HIV-1 subtypes (**Fig. 2f** and **2g**).

**Fig. 2.**
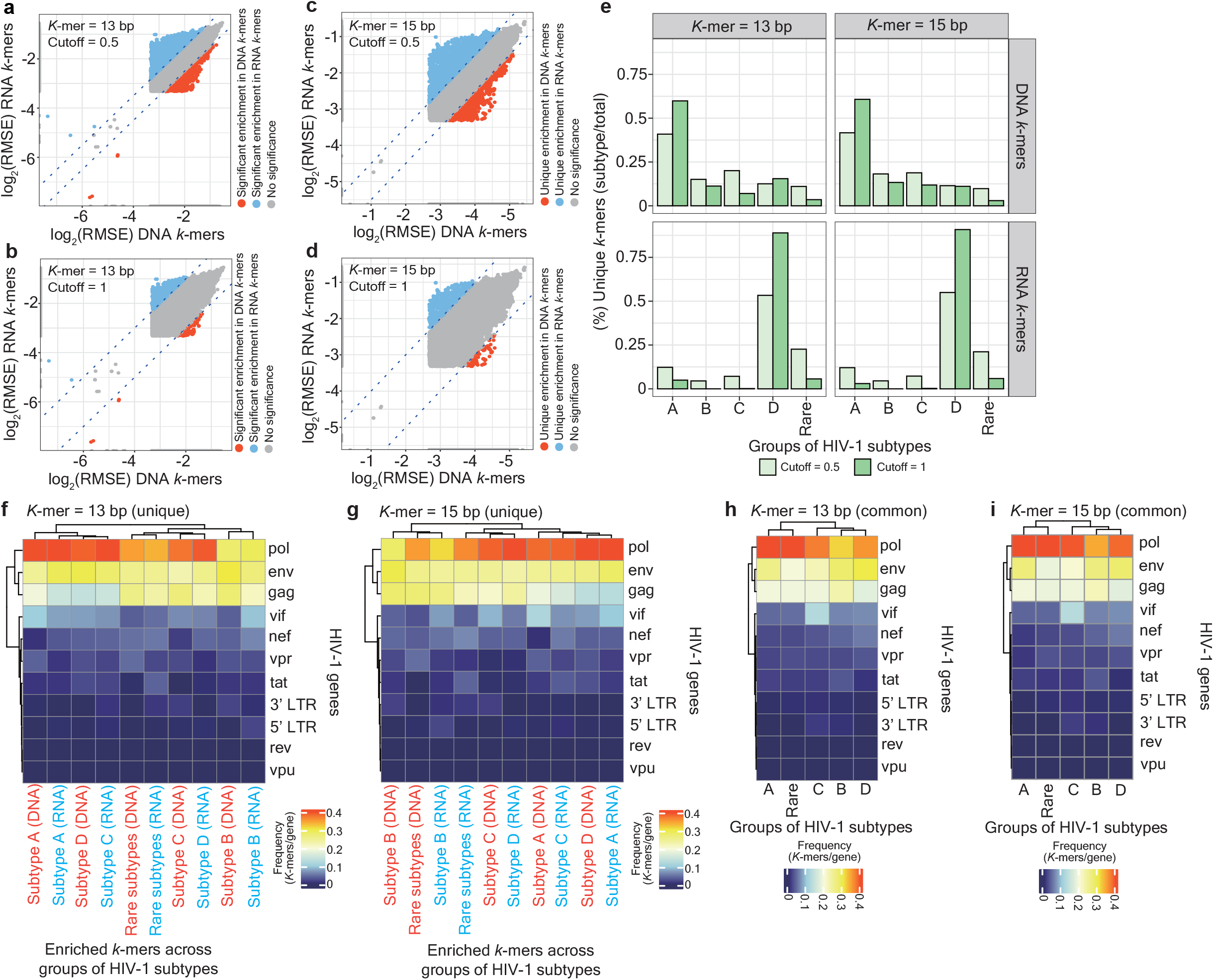
Unique enriched *k*-mers account for the discrepancy of the sequence patterns across HIV-1 subtypes. (**a-d**) Scatter plots representing the selection of unique enriched DNA and RNA *k*-mers in a length of 13 bp (**a** and **b**) and 15 bp (**c** and **d**). Two distinct cutoffs (**a** and **c**, cutoff = 0.5; **b** and **d**, cutoff = 1) based on the ratio of “*k*-mer RMSE” represented on a logarithmic scale were applied. Dots marked in red-orange and light blue represent enriched *k*-mers, which are exclusively present in a pool of enriched DNA and enriched RNA *k*-mers, respectively; dots marked in grey represent enriched *k*-mers that do not pass the selection threshold. (**e**) Bar charts representing the percentage of unique DNA and RNA *k*-mers in two distinct cutoffs (cutoff = 0.5 versus cutoff = 1.0) across different groups of HIV-1 subtypes. Facets at the x-axis separate enriched *k*-mers in a length of 13 bp or 15 bp; facets at the y-axis separate enriched DNA and RNA *k*-mers. Squares marked in light and dark green represent the cutoff threshold at 0.5 and 1.0, respectively. (**f, g**) Clustering heatmaps representing the frequency of unique enriched DNA (written in red beneath the heatmap) and RNA (written in blue beneath the heatmap) enriched *k*-mers aligned to HIV-1 genes (row) in a *k*-mer length of 13 bp (**f**) and 15 bp (**g**) across different groups of HIV-1 subtypes. The color scale depicts the frequency of unique enriched *k*-mers appearing in HIV-1 genes. (**h, i**) Clustering heatmaps representing the frequency of common enriched k-mers in a length of 13 bp (**h**) and 15 bp (**i**) across different groups of HIV-1 subtypes. The color scale depicts the frequency of common enriched *k*-mers appearing in HIV-1 genes.

With respect to identical *k*-mers that are enriched in both DNA and RNA sequences, namely common enriched *k*-mers, separated by different groups of HIV-1 subtypes, a positive correlation in “*k*-mer RMSE” (**Supplementary Fig. S3g**, *k*-mer = 13 bp, *R*^*2*^ = 0.63; **Supplementary Fig. S3h**, *k*-mer = 15 bp, *R*^*2*^ = 0.6), “*k*-mer weight” (**Supplementary Fig. S3i**, *k*-mer = 13 bp; **Supplementary Fig. S3j**, *k*-mer = 15 bp), and “subtype *k*-mer count” (**Supplementary Fig. S3g**, *k*-mer = 13 bp, *R*^*2*^ = 0.89; **Supplementary Fig. S3h**, *k*-mer = 15 bp, *R*^*2*^ = 0.88) were detected. These findings suggest that the minimal contribution to the distinct sequence pattern results from common enriched *k*-mers. No difference in the frequency of enriched unique (**Fig. 2f** and **2g**) and common (**Fig. 2h** and **2i**) *k*-mers that overlap HIV-1 genes was observed.

### “Isolate *k*-mer count” enables the classification and prediction of different groups of HIV-1 subtypes

To evaluate the biological importance of five features associated with enriched *k*-mers, we constructed classifiers based on three approaches: logistic regression (**Fig. 3a**), multinomial logistic regression (**Fig. 3b**), and the neural network (**Fig. 3c**). We observed that a model constructed by “isolate *k*-mer count” surpassed others in the prediction of the origin of enriched *k*-mers (DNA versus RNA) (**Fig. 3a**). Notably, “isolate *k*-mer count” enabled the prediction of the likelihood of different groups of HIV-1 subtypes, with diverse prediction power between enriched DNA and RNA *k*-mers across distinct groups of HIV-1 subtypes (**Fig. 3b**). Such distinct predictive performance may imply the involvement of additional determinants (e.g., inherent genetic and biological variations, and sequence recombinations) in addition to the cause of the difference in the readthrough between DNA and RNA sequences. Prediction performance of “isolate *k*-mer count” in the context of HIV-1 subtypes was also confirmed by a neural network-based model (**Fig. 3c**). The values of AUC (**Fig. 3a**), probability (**Fig. 3b**), and accuracy (**Fig. 3c**) are summarized in **Supplementary Tables S10, S11**, and **S12**, respectively. All these findings signify the importance of the feature “isolate *k*-mer count”. We further computed the Euclidean distance between two isolates based on “isolate *k*-mer count” and observed a clear separation in isolates across subtypes A, B, and C, irrespective of enriched DNA and RNA *k*-mers (**Fig. 3d-3g**). In contrast, isolates in subtypes D and the remaining subtypes were indistinguishable (**Fig. 3d-3g**).

**Fig. 3.**
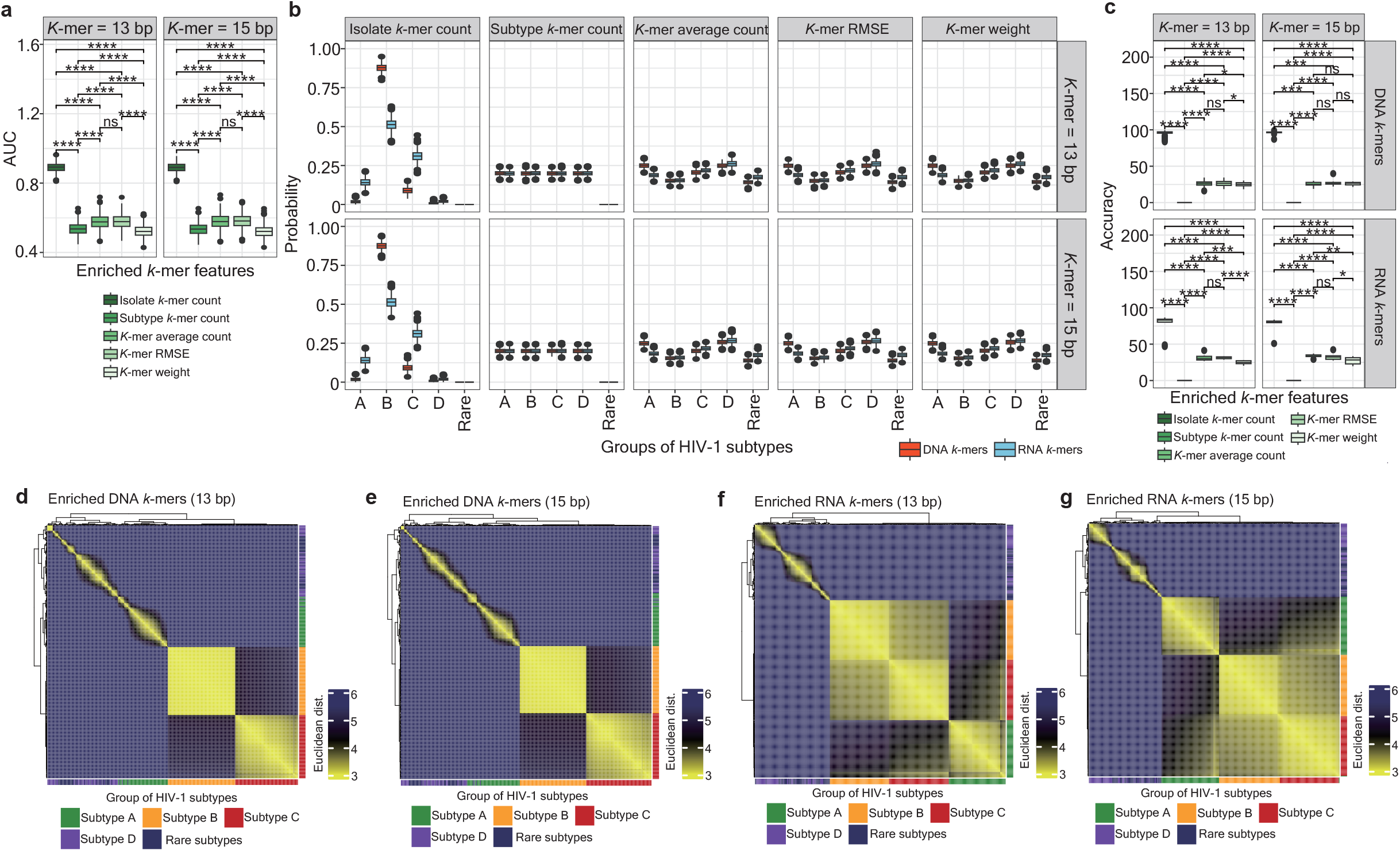
“Isolate *k*-mer count” enables the classification of different groups of HIV-1 subtypes. (**a**) Box plots representing the area under the curve (AUC) of logistic regression models, constructed by distinct enriched *k*-mer-related features, including “isolate *k*-mer count”, “subtype *k*-mer count”, “*k*-mer average count”, “*k*-mer RMSE”, and “*k*-mer weight” (from left to right at the x-axis), to classify the origin of enriched *k*-mers retrieved between DNA and RNA sequences. Each classifier was constructed by 1,000 enriched *k*-mers bootstrapped from the pool of total enriched DNA or RNA *k*-mers, separated between a *k*-mer length equal to 13 bp (the left-hand side panel) and 15 bp (the right-hand side panel). Each classification iteration was repeated 1,000 times for statistical significance. Significance levels are denoted as follows: ns for no significance, \*\*\*\**p* 0.0001. (**b**) Box plots representing the probability of multinomial logistic regression models, constructed by five enriched *k*-mer-related features to classify the origin of enriched *k*-mers retrieved across different groups of HIV-1 subtypes. Each classifier was constructed by 1,000 enriched *k*-mers bootstrapped from the pool of total enriched DNA or RNA *k*-mers, separated between a *k*-mer length equal to 13 bp (top panels) and 15 bp (bottom panels). Each classification iteration was repeated 1,000 times for statistical significance. Facets at the x-axis separate different enriched *k*-mer-related features; facets at the y-axis separate a *k*-mer length between 13 bp and 15 bp. Boxes marked in red-orange and sky blue represent enriched DNA and RNA *k*-mers, respectively. (**c**) Box plots representing the accuracy of simple neural network-based models, constructed by five enriched *k*-mer-related features to verify the predictive performance contributed by individual enriched *k*-mer-related features in the classification of different groups of HIV-1 subtypes, shown in (**b**). Significance levels are denoted as follows: ns for no significance, \**p* 0.05, \*\*\*\**p* 0.0001. (**d-g**) Clustering heatmaps representing the Euclidean distance between two adjacent isolates presenting a topological network constructed by “isolate *k*-mer count” computed from enriched DNA (**d**, *k*-mer = 13 bp; **e**, *k*-mer = 15 bp) and RNA (**f**, *k*-mer = 13 bp; **g**, *k*-mer = 15 bp) *k*-mers across different groups of HIV-1 subtypes. The color scale represents the Euclidean distance between two adjacent isolates. The color code annotation beneath and at the right-hand side of the heatmap denotes the group of HIV-1 subtypes: green, orange, red, purple, and dark blue represent HIV-1 subtype A, B, C, D, and rare subtypes, respectively.

### Intrinsic sequence barriers lie across group M HIV-1 subtypes

We further constructed a pentapartite graph (see **Supporting information** for the definition), representing a network consisting of the five groups of HIV-1 subtypes, and used this graph to illustrate the spatial interactions between isolates (**Supplementary Fig. S4a-S4d**) or enriched *k*-mers (**Supplementary Fig. S4f-S4i**). Overall, *k*-mers that are enriched in the same groups of HIV-1 subtypes tended to connect. In the former scenario, a relatively high assortativity degree was observed in subtypes A, B, and C, whereas a relatively low assortativity degree was present in subtype D and the rare subtypes (**Supplementary Fig. S4e**). In the latter scenario, although a strong tendency between enriched *k*-mers in the same group of HIV-1 subtypes was structured, subsets of *k*-mers enriched in the rare subtypes were observed to link distinct enriched *k*-mer hubs (**Supplementary Fig. S4f-S4i**). A relatively high assortativity degree was observed in subtype B, whereas other subtypes, especially subtype D and the rare subtypes, demonstrated the lowest assortativity degree (**Supplementary Fig. S4j**).

Lastly, we applied the Markov chain Monte Carlo (MCMC) method^52^ and simulated 10 paths of random walks up to 10,000 steps (**Fig. 4a**) in networks constructed by either “isolate *k*-mer count” (**Fig. 4b-4f**) or “subtype *k*-mer count” (real, **Fig. 4g-4k**; labeling mismatch, **Supplementary Fig. S5a-S5e**), respectively. We inspected five different assumptions, in which a random walk initiates in an isolate or an enriched *k*-mer in any HIV-1 subtype group and computed the probability that isolates or enriched *k*-mers in each group of HIV-1 subtypes involve in a path of a random walk (**Fig. 4a**; see **Methods**).

**Fig. 4.**
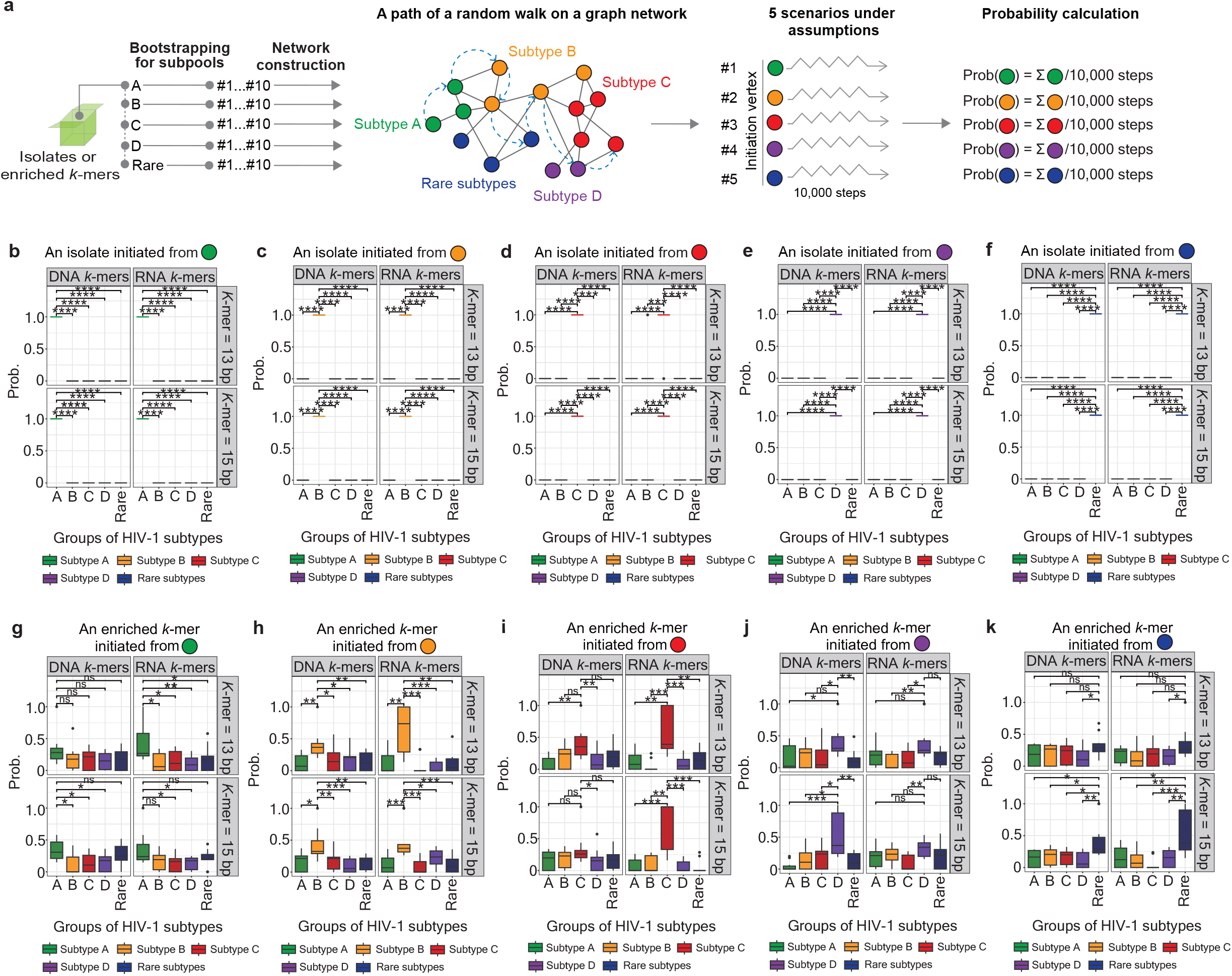
MCMC modeling the intrinsic sequence barrier between HIV-1 DNA and RNA. (**a**) Schematic representation of the strategy of subsampling of enriched *k*-mers, followed by the construction of a pentapartite graph of a topological network. Five scenarios under a framework of assumptions were set up, in which 10 paths of random walks up to 10,000 steps were simulated across distinct network properties. Lastly, the probability of appearing vertices assigned to different groups of HIV-1 subtypes within a path in each scenario was computed. (**b-f**) Box plots representing the probability that vertices assigned to different groups of HIV-1 subtypes appear within a path of each scenario, whereby a random walk initiates a vertex assigned to HIV-1 subtype A (**b**), B (**c**), C (**d**), D (**e**), and rare subtypes (**f**). Networks were constructed by “isolate *k*-mer count”, whereby each vertex represents an individual HIV-1 isolate. Facets at the x-axis separate enriched DNA and RNA *k*-mers; facets at the y-axis separate a *k*-mer length between 13 bp and 15 bp. Boxes marked in green, orange, red, purple, and dark blue represent HIV-1 subtype A, B, C, D, and rare subtypes, respectively. Significance levels are denoted as follows: \*\*\*\**p* 0.0001. (**g-k**) Box plots representing the probability that vertices represent enriched *k*-mers to different groups of HIV-1 subtypes within a path of each scenario, whereby a random walk initiates a vertex assigned to HIV-1 subtype A (**g**), B (**h**), C (**i**), D (**j**), and the rare subtypes (**k**). Networks were constructed by “subtype *k*-mer count”. Facets at the x-axis separate enriched DNA and RNA *k*-mers; facets at the y-axis separate a *k*-mer length between 13 bp and 15 bp. Boxes marked in green, orange, red, purple, and dark blue represent HIV-1 subtype A, B, C, D, and rare subtypes, respectively. Significance levels are denoted as follows: ns for no significance, \**p* 0.05, \*\**p* 0.01, \*\*\**p* 0.001.

In a network constructed by “isolate *k*-mer count”, a superior subtype-specific correlation between initiation of isolates and the highest probability that isolates categorized into the identical group upon the initiation step was observed (**Fig. 4b-4f**), resonating with disconnected graphs (**Supplementary Fig. S4a-S4d**). A similar pattern appeared in networks constructed by “subtype *k*-mer count” (**Fig. 4g-4k**), with more significant discrepancy demonstrated in paths initiating from an *k*-mer enriched in subtype B (**Fig. 4h**) and C (**Fig. 4i**). No difference in labeling mismatch networks across each tested assumption was observed (**Supplementary Fig. S5**). Altogether, these findings verified the presence of intrinsic barriers between genomic sequences across different groups of HIV-1 subtypes.

## Discussion

PORT-EK-v2 has achieved major performance improvements, offering minimal computing time (PORT-EK^47^, 01:01:27; PORT-EK-v2, 00:06:50 [hh:mm:ss]), efficient computing costs, and enhanced analytical capabilities for multigenomic datasets when contrasted with PORT-EK^47^. In other words, PORT-EK-v2 minimizes manual intervention while maintaining high performance accuracy and is readily adaptable to different multigenomic datasets. In addition, the improvement of the enriched *k*-mer mapping strategy—pre-calculated reference genome index—has significantly leveraged the performance, resulting in faster execution, the ability to map recurring *k*-mers, and precise indel handling (detailed in **Supporting information** with the pseudocode provided). Overall, PORT-EK-v2 is an efficient and streamlined pipeline designed to immediately provide *k*-mer frequency enrichment statistics in a readable form when dealing with a high volume of small multigenomic datasets. Optimal parameterization in PORT-EK-v2 is detailed in **Supporting information** and **Supplementary Table S1**. Using PORT-EK-v2, we analyzed multiple HIV-1 DNA and RNA sequences and provided several fresh insights based on findings presented in this work.

Firstly, distinct behaviours between unique and common enriched *k*-mers (**Fig. 2** and **Supplementary Fig. S3**) suggest the presence of subgenomic locations in the HIV-1 genomic composition. Unique enriched *k*-mer frequencies are most likely the main determinants governing intrinsic differences across different HIV-1 subtypes. The phenomenon is remarkably observed in HIV-1 subtype B, when DNA sequence is analyzed (**Supplementary Fig. S3b** and **S3d**). For at least three decades, the epidemic in the Western World has been dominated by subtype B infections^53^. At present, subtype B remains the most disseminated variant and is assumed to be the causative agent in approximately 11% of all cases of HIV worldwide^54^. Relative to other group M HIV-1 subtypes, a higher genomic diversity measured based on enriched *k*-mer frequency in subtype B may be reflected by its global spread, thereby accelerating the evolution of its genomic composition (e.g., accumulation of mutations or deletions, and so on). This assumption could be underpinned by the observation of an outlier of enriched *k*-mers (**Fig. 1a** and **1b**). Further investigation of the evolutionary trajectory of subtype B at the sequence or nucleotide levels will be required to gain a more insightful understanding of its genomic diversity at a larger time scale.

Secondly, an indistinguishable pattern between enriched DNA and RNA *k*-mers mapped to HIV-1 genes may infer insufficiency of resolution for the classification of the sequence property at the genic level (**Fig. 2f-2i**). This also suggests that future taxonomies of HIV-1 subtypes and the phylogenetic analysis should be manifested in a higher-tiered order at the levels of short pieces of genomic sequences or nucleotides to discover more profound diversities of HIV-1 subtypes and recombinant forms. It is important to take into account the inconsistency between HIV-1 DNA and RNA appearing in the same group of HIV-1 subtypes (**Fig. 2e**), which may lead to different biological interpretations. In addition, one should also seek informative features from non-coding regions, given that the majority of variants identified by genome-wide association studies of complex phenotypes are non-coding^55^.

Thirdly, the consistent gap present (ranging from 8.3 kb to 8.6 kb) across all HIV-1 subtypes in both enriched DNA and RNA *k*-mer profiles (**Supplementary Fig. S3e** and **S3f**) indicates that nucleotide compositions in this gap are highly varied across HIV-1 subtypes. Of note, no conserved *k*-mers were detected in this region.

Intriguingly, this region overlaps with the second exon of *tat*-*rev* (8,433-8,459 bp) and is in the vicinity of the HIV *rev* response element (RRE, 7,832-7,851 bp)^56-58^. HIV-1 Tat-Rev are essential regulatory proteins that drive HIV-1 gene expression and replication^59^ and govern the switch from early-phase to late-phase viral replication, ensuring the delicate balance between early and late infection^60^. Biological consequences of this consensus genomic region will require further investigation.

Fourthly, our findings that relatively high “isolate *k*-mer count” in the rare subtypes (**Supplementary Fig. S1h**) and an independent subset in the group of the rare subtypes in phylogenetic trees constructed using enriched RNA *k*-mers (**Fig. 3j** and **3k**) unveil the genomic heterogeneity within the rare HIV-1 subtypes, supported by the fact that many of these rare subtypes are more frequently found in recombinant forms^28,61– 65^. Although subtypes C and B are the most prevalent globally^53,54^, the rare subtypes, including F, H, J, K, and L, which are individually responsible for less than 1% of HIV-1 infections worldwide, have been reported to play relevant roles in viral evolution, global persistence, recombination, and drug resistance as well^66,67^. In this circumstance, a more sophisticated and sensitive method that enables the detection of the genomic discrepancy at the molecular level will be essential to tackle ongoing viral outbreaks, pandemic preparedness, and to respond to ongoing and emerging threats.

Altogether, in this work, we propose that distinct sequence patterns between HIV-1 DNA and RNA across group M HIV-1 subtypes exist based on enriched *k*-mers retrieved by PORT-EK-v2, and were verified by different downstream analyses. Results from machine learning-based modeling have verified that the enriched *k*-mer-associated feature, “isolate *k*-mer count”, enables the classification of HIV-1 DNA and RNA genomic sequences (**Fig. 3a**) and the discrimination of different groups of HIV-1 subtypes (**Fig. 3b** and **3c**). Such intrinsic differences were further confirmed by the calculation of Euclidean distances (**Fig. 3d-3g**). Moving forward, the observations based on the graph-based analysis (**Supplementary Fig. S4**) and MCMC simulations (**Fig. 4**) have suggested that this intrinsic difference can also exist at the global level of network organization. Nevertheless, observations based on these different analyses lead us to hypothesize that the presence of intrinsic differences in HIV-1 DNA and RNA sequences across subtypes can be unveiled based on enriched *k*-mers.

### Limitations of this study

One of the major limitations is that recombinant subtypes have not been included in this study, like the circulating recombinant form (CRF) 01_AE, which is one of the most prevalent HIV-1 subtypes, together with subtypes C, A, and B^68^. Given the complexity of the genomic composition of CRF01_AE, a putative recombinant between subtypes A and E^69^, and the uncertainty of the origin of its parental subtype E^70,71^, only the non-recombinant genomic sequences from group M HIV-1 subtypes were applied in this work, enabling a better examination of the feasibility of PORT-EK-v2 and the functionality of enriched *k*-mers. It is also important to note that due to the prevalence of HIV-1 subtype B in Latin America and the Caribbean^72,73^, where viruses later spread to North America, as well as in Europe^74^, a higher volume of subtype B genomic data is available in the Los Alamos HIV Sequence Database compared to other subtypes. While we cannot currently eliminate the possibility of a bias caused by the large volume of subtype B genomic sequences, which could impact result interpretation, our approach to finding *k*-mer frequency showed no obvious link between input data size and enriched *k*-mer counts across different groups of HIV-1 subtypes. Nevertheless, a more meticulous review of artefactual *k*-mers (e.g., those resulting from a bias of input data size or sequencing error, and so on) together with other techniques will be expected to increase the reliability of the extracted *k*-mers. In addition, mutations emerging on enriched *k*-mers have not been investigated yet. Future research focus should be placed on closer investigation of the functional roles (e.g., identification of drug-resistant mutations across distinct HIV-1 subtypes) of *k*-mers between DNA and RNA sequences in clinical management, enabling a better application in the context of HIV-1 molecular epidemiology associated with precise genotypic HIV-1 drug resistance testing^75,76^, and genotypic therapeutics. At present, PORT-EK-v2 has only been examined using multigenomic sequences from HIV-1 subtypes. In the future, it should also be tested using multigenomic sequences from a diverse array of species to determine its viability for a wider spectrum of organisms. It is also crucial to restate that PORT-EK-v2 was initially tailored for a high volume of small genomes and may not be ideal for large-scale genomic data. Nevertheless, PORT-EK-v2 may have the potential to serve as one of the novel molecular epidemiology approaches applied to the future investigation of strategies for curbing HIV transmission, preventing resistance, and improving the understanding of HIV pathogenesis and HIV-1 genomic surveillance, enabling the identification of new and emerging subtypes.

## Methods

### Data acquisition and quality control of retrieved sequences

A total of 10,015 and 5,494 HIV-1 DNA and RNA genomic sequences across different subtypes within the group M HIV-1 were retrieved from the Los Alamos HIV Sequence Database (https://www.hiv.lanl.gov/). Only complete genomic sequences were included in this study. Analyzed sequences and corresponding exact IDs are archived at GitHub (https://github.com/Quantitative-Virology-Research-Group/PORT-EK-version-2/tree/main/Input.HIV.sequence). 2 and 4 HIV-1 DNA and RNA sequences, which cannot be aligned to the reference genome (HIV-1 HXB2, complete genome K03455.1, curated in the National Center for Biotechnology Information database), provided in the Los Alamos HIV Sequence Database were discarded.

### The principle of the Pathogen Origin Recognition Tool using Enriched *K*-mers version 2 (PORT-EK-v2)

PORT-EK-v2 is the upgraded version of PORT-EK^47^, a pipeline that compares sequences from two different multigenomic datasets and identifies over-represented *k-*mers in respective datasets. The principle of *k*-mer-based approaches is to count the number of distinct substrings of length *k* in a string S^77^, or a set of strings, where *k* is a positive integer representing a unique mark in an indicated string S^77^. In this study, we computed five features of enriched *k*-mers, including “*k*-mer average count”, “*k*-mer root mean square deviation (RMSE)”, “*k*-mer weight”, “subtype *k*-mer count”, and “isolate *k*-mer count”, for better characteristics of enriched *k*-mer. Methodology for the calculation of each feature is detailed in **Supporting information** and the following section, wherever applicable. The Python pipeline and related R scripts for downstream analyses are available at GitHub https://github.com/Quantitative-Virology-Research-Group/PORT-EK-version-2. It is worth noting that we optimized the PORT-EK-v2 parameterization—the number of parameters has been reduced from 6, including *k, c, m, min*_*RESM*_, *f*_*c*_, and *l*_*map*_, to 4, including *k, d* (which replaced *m*), *max_mem*, and *min_freq* (see **Supporting information** for the definition of each parameter). Details of each step involved in PORT-EK-v2 are described in **Supporting information**.

### PORT-EK-v2 benchmark versus PORT-EK

Benchmark runs were performed on a desktop PC with AMD Ryzen 7 7700X 8-Core CPU and 32 GB of RAM, running Ubuntu 24.04.01 LTS WSL for Windows 11. Benchmark results are released in **Tables 1** and **2**.

### Computation of the feature of enriched *k*-mers

Five features, including “*k*-mer average count”, “*k*-mer RMSE”, “*k*-mer weight”, “subtype *k*-mer count”, and “isolate *k*-mer count”, were included in this study for a better characterization of enriched *k*-mers. The rationale of computing “*k*-mer average count” and “*k*-mer RMSE” has been detailed in **Supporting information**, together with the rationale of PORT-EK-v2. *K-mer weight*: This attribute was designated to unveil the composition of enriched *k*-mers. Here, we applied ordinal encoding, which presents baseline encoding, to represent each of the four nucleotides (A, T, C, and G) as a number between 0 and 1. We first utilized the previous setting of ordinal encoding from the study of Wade et al. (2024)^43^, whereby A is encoded as 0.25, T as 0.50, C as 0.75, and G as 1.00, and summed the weights for individual enriched *k*-mers. The “*k*-mer weight” is defined by the Equation below (**Fig. 1i**).

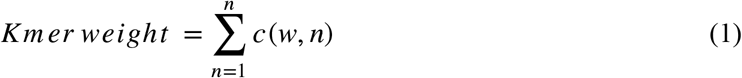

where *n* is the index of each nucleotide over a string of an enriched *k*-mer sequence, and *c* (*w, n*) represents the assigned ordinal encoding value *w* in a specific nucleotide *n*. The computing code is written in R and available at GitHub (https://github.com/Quantitative-Virology-Research-Group/PORT-EK-version-2/blob/main/Analysis/Main.figures/Fig_1f_Kmer.weight.R). Alterations of ordinal encoding assigned to four nucleotides, as controls, were performed and illustrated in **Supplementary Fig. S2**.

#### Subtype k-mer count

We aggregated the count of individual *k*-mers across all isolates present in the same group of the HIV-1 subtype; values were then normalized by the total number of enriched *k*-mers present in the same group of the HIV-1 subtype. The “subtype *k*-mer count” is defined by the Equation below (**Fig. 1i**).

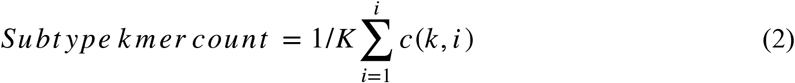

where ***K*** is the total number of enriched *k*-mers in an indicated group, *i* is the index over all isolates in the indicated group, and *c*(*k*, *i*) represents the count of enriched *k*-mer *k* in an isolate *i*. The computing code is written in R and available at GitHub (https://github.com/Quantitative-Virology-Research-Group/PORT-EK-version-2/blob/main/Analysis/Main.figures/Fig_1g_subtype.kmer.count.R).

#### Isolate k-mer count

We aggregated *k*-mer counts by HIV-1 isolates present in the same group of HIV-1 subtype. The sum was normalized by the total number of isolates present in the same group of the HIV-1 subtype. The “isolate *k*-mer count” is defined by the Equation below (**Fig. 1i**).

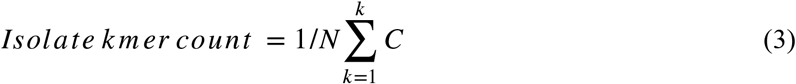

where *N* is the total number of isolates in an indicated group, and *C* is the summation of all enriched *k*-mer counts (to the *k*-th enriched *k*-mer) in an individual isolate. The computing code is written in R and available at GitHub (https://github.com/Quantitative-Virology-Research-Group/PORT-EK-version-2/blob/main/Analysis/Main.figures/Fig_1h_isolate.kmer.count.R).

### Construction of prediction models

#### Logistic regression

To examine the prediction power in the classification of the origin of enriched *k*-mers (DNA versus RNA *k*-mers), 1,000 random enriched *k*-mers from each individual pool of enriched *k*-mers were bootstrapped with replacement and divided into a training set (80% of the dataset) and a testing set (20% of the dataset) for logistic regression running on R. The logistic regression model was fitted using the function glm() with the argument family specified as “na.exclude” and ‘‘binomial’’ in the R package ‘‘stats’’ (https://www.r-project.org/). Classifiers associated with the five mentioned features (i.e., “*k*-mer average count”, “*k*-mer RMSE”, “*k*-mer weight”, “subtype *k*-mer count”, and “isolate *k*-mer count”) were independently constructed; the “response” (i.e., the origin of enriched *k*-mers), coupled with a corresponding *k*-mer feature, was thus included in an object of class “formula”, independently. Receiver operating characteristic (ROC) and the area under the curve (AUC) were calculated using the functions multiclass.roc() and auc() in the R package ‘‘pROC’’^78^, respectively. Given that there is no great variety of the input data for modeling, we have chosen ROC-AUC for evaluating a model’s overall ranking ability across all thresholds. The whole procedure was repeated 1,000 times for statistical robustness. The computing code is written in R and available at GitHub (https://github.com/Quantitative-Virology-Research-Group/PORT-EK-version-2/blob/main/Analysis/Main.figures/Fig_1g_Kmer.count.isolate.R).

#### Multinomial logistic regression

To examine the prediction power in the classification of the distinct groups of HIV-1 subtypes based on enriched *k*-mers, 1,000 random enriched *k*-mers from each individual pool of enriched *k*-mers were bootstrapped with replacement and divided into a training set (80% of the dataset) and a testing set (20% of the dataset) for constructing multinomial logistic regression-based models associated with five features, running on R. The multinomial logistic regression model was fitted using the function multinom() in the R package ‘‘nnet’’^79^. Classifiers associated with the five mentioned features were independently constructed. The predicted probability that enriched *k*-mers falling into distinct groups of HIV-1 subtypes, including subtypes A, B, C, D, and the rare subtypes, coupled with a corresponding *k*-mer feature, was computed using the function predict() with the argument family specified as type = “probs” in the R package ‘‘stats’’ (https://www.r-project.org/). Given that there is no great variety of input data for modeling, we have calculated the probability for evaluating the model’s confidence level. Output probabilities were conserved in a dataframe. The whole procedure was repeated 1,000 times for statistical robustness. The computing code is written in R and available at GitHub (https://github.com/Quantitative-Virology-Research-Group/PORT-EK-version-2/blob/main/Analysis/Main.figures/Fig_3b_Kmer.multinomial.LG.subtypes.R).

#### Simple neural network

We performed simple neural network modeling to verify the prediction result obtained from multinomial logistic regression-based models. 1,000 random enriched *k*-mers from each individual pool of enriched *k*-mers were bootstrapped with replacement and divided into a training set (80% of the dataset) and a testing set (20% of the dataset) for the performance of simple neural network running on R. The model was fitted using the function neuralnet() in the R package ‘‘neuralnet’’^80^. Classifiers associated with the five mentioned features were independently constructed. The whole procedure was repeated 10 times for statistical robustness. The computing code is written in R and available at GitHub (https://github.com/Quantitative-Virology-Research-Group/PORT-EK-version-2/blob/main/Analysis/Main.figures/Fig_3c_Kmer.NN.R).

It is important to note that the input data representing “isolate *k*-mer count” is a HIV-1 isolate-based matrix. While their balance was first managed during the PORT-EK-v2 processing phase for obtaining enriched *k*-mers, the inherent bias resulting from the unequal distribution of isolates among HIV-1 subtypes, restricted by current data availability, must still be acknowledged.

### Calculation of the Euclidean distance between isolates

Euclidean distance is chosen based on the consideration that the magnitude (quantity) of the “isolate *k*-mer count” matters. Given that the length of a *k*-mer has been restricted and the network has been resampled to an identical size of 100 isolates for each group of HIV-1 subtypes, their influence is deemed to be negligible. To reduce computing cost, we bootstrapped 100 enriched *k*-mers and repeated 10 times to create 10 subpools of enriched *k*-mers in each group of HIV-1 subtypes. In each subpool of enriched *k*-mers, an edge weight between two isolates is calculated by the sum of “isolate *k*-mer count” normalized by the total number of isolates in the corresponding group of HIV-1 subtypes. The edge weight is defined by the Equation below.

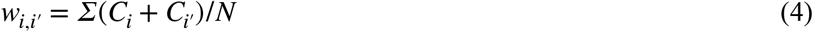

where 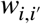 is the edge weight between isolate *i* and isolate, 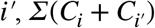 is the sum of the *k*-mer counts for isolate *i* and isolate *i*′, respectively, and *N* is the total number of isolates in an indicated group.

The computed edge weight was assigned to individual isolates and was used to compute a matrix of the Euclidean distance, together with the parameter “isolate *k*-mer count” at an individual isolate level using the function dist() with the argument family specified as method = “euclidean” in the R package ‘‘stats’’ (https://www.r-project.org/). The clustering heatmap was visualized using the R package “ComplexHeatmap”^81,82^. The computing code is written in R and available at GitHub (https://github.com/Quantitative-Virology-Research-Group/PORT-EK-version-2/blob/main/Analysis/Main.figures/Fig_3e_3h_Euclidean.dist.heatmap.R).

### Phylogenetic tree analysis

Unrooted phylogenetic trees were constructed from enriched *k*-mer count matrices using a distance-based approach. Optionally, the user can randomly subsample the dataset to a specified number of genomes, either uniformly or with balanced representation of multiple genomes. In this study, all datasets were downsampled to 50 genomes of each HIV-1 subtype, 250 genomes per tree. Pairwise Euclidean distances between all genomes’ *k*-mer count vectors were calculated using *pdist* function from *scipy*.*spatial*.*distance* l ibrary. A distance matrix was then created with *Biopython ‘s Phylo*.*TreeConstruction*.*DistanceMatrix* class. Finally, a phylogenetic tree was inferred from this matrix with *Biopython’s Phylo*.*TreeConstruction*.*DistanceTreeConstructor* class, using the neighbor-joining (nj) method. The resulting tree object was written to Newick file format (.nwk) using the Phylo._io.write function from *Biopython*. The file was then read with the function read.tree() and visualized with the function ggtree() in the R package “ggtree”^83^ with the %<+% operator to connect the annotation files corresponding to each phylogenetic tree.

### Markov chain Monte Carlo modeling analysis

A concatenation of networks were constructed by either “isolate *k*-mer count” or “subtype *k*-mer count” across different groups of HIV-1 subtypes for simulation (see **Supporting information**). 10 networks constructed by presampling 100 isolates were computed; the readouts of pairwise correlation matrices were saved as a list of data frames. A list of data frames was then expanded by generating all possible pairs of vertices, representing either “isolate *k*-mer count” or “group *k*-mer count” using the function complete() in the R package “tidyr” (https://tidyr.tidyverse.org). Any pair of vertices with the edge coefficient (i.e., summation of either “isolate *k*-mer count” or “group *k*-mer count” between two adjacent vertices) equal to zero was removed. Moving forward, a list of expanded data frames was reshaped from long to wide format using the function dcast() with the option “fun.aggregate = sum” in the R package “maditr” (https://github.com/gdemin/maditr). We further coordinated each vertex present in a list of expanded data frames with corresponding pairwise correlation matrices, enabling the reassignment of each vertex to respective groups of HIV-1 subtypes.

Five scenarios under the assumption, in which a random walk commences with an isolate or an enriched *k*-mer assigned to different groups of HIV-1 subtypes, were established. Based on the coordinate between each vertex and the corresponding group of HIV-1 subtypes, we forced a random walk beginning with the first appearing vertex in different groups of HIV-1 subtypes and completing within a total of 10,000 steps. In principle, in each step we first sought and listed all neighbor vertex that possess the edge coefficient greater than zero from the current vertex, and randomly selected one neighbor vertex from this list with equal probability using the R built-in function sample(). Lastly, we computed the probability of appearing vertices assigned to different groups of HIV-1 subtypes within a path in each scenario. The computing code is written in R and available at GitHub (https://github.com/Quantitative-Virology-Research-Group/PORT-EK-version-2/blob/main/Analysis/Main.figures/Fig_4b_4f_random.walk.isolate.R and https://github.com/Quantitative-Virology-Research-Group/PORT-EK-version-2/blob/main/Analysis/Main.figures/Fig_4g_4k_random.walk.subtype.R).

## Data visualization

All plots were visualized using R (version 4.4.1) with the packages “ggplot2” (https://ggplot2.tidyverse.org), “ ComplexHeatmap”^81,82^, “ggtree”^83^, “treeio”^84^, and “igraph”^85^.

## Statistics

All statistical tests were performed with R with default options. Details are provided where appropriate in the main text.

## Data availability

The final output of enriched DNA and RNA *k*-mers can be referred to **Supplementary Table S2-S5**. Enriched DNA and RNA *k*-mer counts at the coordinate of isolates can be referred to **Supplementary Table S6-S9** (archived at Zenodo DOI:10.5281/zenodo.18757144).

## Code availability

The PORT-EK-v2 code, the instruction of installation and execution of the PORT-EK-v2 pipeline, and all codes for downstream analyses performed in this work are available at GitHub (https://github.com/Quantitative-Virology-Research-Group/PORT-EK-version-2).

## Supplementary Data statement

Supplementary Tables S1-S12.

Tables S2-S9. Excel files containing additional data too large to fit in a PDF and are archived at Zenodo (DOI:10.5281/zenodo.18757144).

## Acknowledgements

The study was partially supported by the IMPRESS-U program, funded by the Narodowe Centrum Nauki (NCN, Poland; grant UMO-2023/05/Y/ST6/00263) and the National Science Foundation (NSF, USA; grant OISE-2412914).

## Author Contributions Statement

Conceptualization: HCC

Methodology: JW and HCC

Software: JW and HCC

Formal analysis: JW and HCC

Investigation, JW, SK, and HCC

Resources: HCC

Data curation: JW and HCC

Writing of original draft manuscript: JW and HCC

Writing, manuscript review and editing: JW, SK, MP, AKP, PS, AK, SY, and HCC

Visualization: JW and HCC

Supervision: HCC

Project administration: AK, SY, and HCC

Funding acquisition: AK, SY, and HCC

