## Supporting information for "Identification of different intrinsic sequence patterns between HIV-1 DNA and RNA across subtypes using the *k*-mer-based approach"

### Supplementary information

**The rationale of the design of the Pathogen Origin Recognition Tool using Enriched *K*-mers version 2 (PORT-EK-v2).** While using PORT-EK-v2, input files should be in the fasta format with headers formatted as in NCBI, GISAID, or any other format not containing forward slashes "/". All sequences in one input file should correspond to the same multigenome, and there should be one file per multigenome. The names of the input fasta files, the names of their corresponding multigenomes and header formats, as well as the reference sequence file name and optional list of reference gene intervals must be provided in a config.yaml file for each dataset. A template of the configuration file and an example of a complete config file are provided with the software. PORT-EK-v2 allows the comparison of an arbitrary number of independent sources of multigenomes, either in all possible pairs ("all-vs-all" or "ava" mode) or one control multigenome against all others ("one-vs-rest" or "ovr" mode). In this study, each dataset consists of multiple genomes isolated from five different groups across HIV-1 subtypes, and "all-vs-all" mode was used. Three tunable parameters, the *k*-mer length *k,* maximum *k*-mer count matrix size *max_mem*, minimum *k*-mer frequency *min_freq* (used only when count matrix size would surpass *max_mem*), and allowed mapping edit distance *d* are used. Of note, many of the arbitrary parameters used by PORT-EK[47] that required adjustment whenever a new genomic dataset was applied, including conservation threshold *c*, allowed rare *k*-mer mismatches *m*, minimum root mean square error $min_{RMSE}$, and allowed mapping offset $l_{map}$, are no longer required in PORT-EK-v2. Moreover, PORT-EK-v2 automatically suggests the best *k* value between requested *min_k* and *max_k* (default 5 and 31). A systematic comparison of parameters used between PORT-EK[47] and PORT-EK-v2 is summarized in **Supplementary Table S1**. The details of the parameters and their settings in PORT-EK-v2 are described in the following section. Recorded running times and peak memory usage for benchmark data sets are presented in **Table 1**. PORT-EK-v2 is written in Python 3.12.3. It uses Biopython 1.85[86], matplotlib 3.10.7[87], numpy 2.3.4[88], pandas 2.3.3[89], scikit-learn 1.7.2 (Pedregosa *et al.* Scikit-learn: Machine Learning in Python, JMLR 12, pp. 2825-2830, 2011), scipy 1.16.2[90], and seaborn 0.13.2[91] libraries.

*K-mer extraction, count matrix, and descriptive statistics used in PORT-EK-v2*

We first extracted *k*-mers of length *k* with overlapping sequences using a sliding window, moving every nucleotide and placing them in hash table files. Any *k*-mer sequences containing ambiguous nucleotides were discarded. The sequences and the list of first-nucleotide positions were used as key-value pairs. HIV-1 subtypes were appended to corresponding hash table file names. For brevity, we termed these files *k*-mer position indices, or simply indices in the rest of this work. Of note, the *k*-mer sequences are encoded as integers, and the indices are stored as binary files. This reduces the required disk space and increases file I/O speed compared with PORT-EK[47], which stored the sequences in text format as JSON files. Simultaneously with *k*-mer extraction, two statistics were calculated. Frequency of each *k*-mer for each subtype, $f_i$, was calculated based on the ratio between the total number of genomes containing a particular *k*-mer and the total number of genomes in the respective subtype. The average count of each *k*-mer for each HIV-1 subtype, $n_i$, was calculated as the arithmetic mean.

Based on the assumption that *k*-mers containing meaningful information should not be very rare or extremely common[92], we discard *k*-mers present in every genome or only single genomes. If, however, the resulting *k*-mer count matrix would exceed *max_mem* GB in size, we additionally discard *k*-mers with frequency $f_i$ lower than *min_freq* in every subtype. Higher settings of *max_mem* allow the pipeline to capture rarer variants that may still be meaningful at the cost of increased computation time, memory usage, and difficulty of interpretation. Next, the indices were read to construct a *k*-mer count matrix containing the count of every remaining *k*-mer in every genome, with rows indicating the *k*-mer sequence and columns indicating genomes.

This approach of calculating basic *k*-mer statistics $f_i$ and $n_i$ and using them to filter out rare *k*-mers before the construction of the count matrix, significantly reduced the computing time and memory requirements of PORT-EK-v2 compared to PORT-EK[47] (**Table 1**), which performed similar filtering after the construction of the full *k*-mer count matrix.

We further computed the following three statistics on common *k*-mers:
(i) the difference in the average count of each *k*-mer between each paired subtype in "all-vs-all" mode or between the control subtype and all other subtypes in "one-vs-rest" mode: $\Delta n_1 = n_1 - n_2$, $\Delta n_2 = n_1 - n_3$, and so on.
(ii) the root mean square error (RMSE) of said changes:

$$RMSE = \sqrt{\frac{1}{s-1}\sum_{i=1}^{s}(\Delta n_i)^2} \quad (5)$$

, where *s* is the total number of subtypes.

The difference in the average count of each *k*-mer was used throughout the whole pipeline as a metric of enrichment, indicating a *k*-mer with a significant enrichment in the average count in one subtype versus other subtypes. In this work, the positive and negative value of the difference denotes the *k*-mer enrichment in one subtype in particular pairwise comparisons. The RMSE captures the enrichment strength in the whole dataset. The larger the RMSE is, the more pronounced the difference between the two compared HIV-1 subtypes is. Finally, the statistical significance was tested for each pairwise comparison using the Mann-Whitney U test. The differences with *p*-values less than 0.01 in all pairwise comparisons were deemed significant. No multiple hypothesis testing correction was performed in this work; however, PORT-EK-v2 is able to optionally apply Benjamini-Yekutieli false discovery rate control[93] when run with *--fdr* argument.

*Determination of the optimal k value*

To determine the optimal *k* value, *k*-mers of length 5 to 31 were extracted as described above, and the average length of the input sequences, *L,* was calculated. Then, *P*, the expected percentage of unique *k*-mers in a random sequence of length *L* was calculated for all tested *k* values:

$$P = e^{-\frac{(L-k+1)^2}{2*4^k}} \quad (6)$$

*P* increases quickly for small values of *k* and plateaus at values near 100% for larger values of *k,* forming a scree plot. Optimal *k* was selected as the onset of the plateau as determined by the Kneedle algorithm[94]. This allows for the selection of the lowest *k* value that still can adequately capture the diversity of the dataset, saving resource usage. PORT-EK-v2 automatically computes the optimal length of a *k*-mer corresponding to its input data. Based on the input HIV-1 DNA and RNA sequences conducted in this work, a *k* value of 15 was selected as optimal. We also tested *k* values of 13 and 17 and found no significant differences between these three *k* values.

*Enriched k-mer identification*

Enriched *k*-mers were retrieved from a pool of common *k*-mers based on the three mentioned statistical strategies. Only *k*-mers with *p*-values less than 0.01 were retained. The parameter $\Delta n_i$ was used to determine in which HIV-1 subtypes *k*-mers were enriched. Finally, *k*-mers that have an RMSE less than 0.1 were discarded. The rest of the *k*-mers were considered to manifest significant enrichment and were assigned to the corresponding hosts. Additionally, *k*-mers present exclusively in one multigenome were deemed "exclusive" and this property was noted in a separate column. The enriched *k*-mer matrix was then transposed, as genome IDs were listed in rows and *k*-mers in columns. HIV-1 subtypes were assigned to enriched *k*-mers shown in columns. Of note, *k*-mers present in more than 90% of all genomes in every multi-genome were deemed conserved *k*-mers and retained regardless of *p*-values or RMSE. Additionally, 2-dimensional PCA decomposition of the *k*-mer counts for all genomes is performed using *scikit-learn.decomposition.PCA* class and scatter plots presenting the results with genomes color-coded by multigenome are created using *seaborn.scatterplot* function, with the variance ratio explained by PC1 and PC2 added to axis labels.

*Enriched k-mers mapping*

The enriched *k*-mers identified in the previous step were then mapped to the HXB2 HIV-1 reference genome with the following algorithm: For a given *k* and maximum allowed number of mismatches *d*, an index of the reference genome was constructed first. At every reference position, the exact *k*-mer was extracted and its 1-based starting position stored. Next, for each *k*-mer, all possible ambiguous *k*-mer patterns representing up to *d* mismatches were generated. This included substitution-like patterns generated by replacing combinations of positions with the wildcard base "N", and simple indel-like patterns generated by inserting stretches of "N" or, when possible, deleting bases within the *k*-mer and appending downstream reference bases. These ambiguous patterns were grouped by the number of mismatches from 0 to *d* and stored in a dictionary. The resulting index was saved to disk as a binary .pkl file and can be reused in subsequent runs. A maximum allowed number of mismatches *d* of 3 was used for all datasets in this work.

After the reference genome index was complete, each enriched *k*-mer was converted into its own set of ambiguous patterns for all numbers of mismatches from 0 to *d*, using the same substitution and indel generation rules. For each number of mismatches, the precomputed index was queried with these patterns and collected all matching reference positions. All corresponding reference positions with the smallest number of mismatches were stored as the mapped positions of the *k*-mer. Then each *k*-mer was mapped to one or more genes by checking whether its mapped positions fall within gene intervals provided in the configuration file. Finally, a per-base coverage matrix over the reference genome was constructed—for each enriched *k*-mer, and for each mapped start position, all reference positions covered by that *k*-mer are incremented in multigenome-specific coverage tracks according to the *k*-mer's assigned multigenome (and in a conserved track for *k*-mers classified as conserved). Two types of output tables were written as the pipeline outputs: (i) a mapping table listing *k*-mers, their mapped reference positions, mismatch number, multigenome, exclusivity, and gene assignment, and (ii) a genome-wide coverage table summarizing enriched *k*-mer coverage per reference nucleotide position and multi-genome.

This approach of matching ambiguous *k*-mer patterns to a pre-calculated reference genome index offers several advantages over the regular expression-based method used in PORTK-EK[47]. First, it has much better performance—With 5,063 genomes in the RNA dataset, PORT-EK-v2 completed the mapping in 30 seconds, while PORT-EK[47] took about 3 minutes. Second, it allows for mapping of *k*-mers that appear more than once per genome, while PORT-EK[47] was limited to mapping unique *k*-mers. Finally, PORT-EK-v2 offers a proficient performance in handling indels, which in PORT-EK[47] could be confused by multiple substitutions. Of note, conserved non-enriched *k*-mers are detectable and can be mapped based on the HIV-1 HXB2 genome. Positions of those *k*-mers are provided together with final enriched *k*-mer outputs using PORT-EK-v2.

The algorithm is illustrated by the pseudocode below:

```
STEP 1 — Index the reference sequence
  IF index file for (ref_seq_name, k, max_n_mismatch) does not exist on disk THEN
    FOR each position in ref_seq DO
      Extract exact k-mer at position → store in index at distance 0
      FOR n_mismatches = 1 TO max_n_mismatch DO
        Generate all substitution variants (replace n positions with "N")
        Generate all insertion/deletion variants of length n
        Store each variant → position mapping in index at distance n
      END FOR
    END FOR
    Write index to disk
  ELSE
    Load index from disk
  END IF

STEP 2 — Map each enriched k-mer to the reference
  FOR each kmer in enriched k-mers table DO
    FOR n_mismatch = 0 TO max_n_mismatch DO
      Generate substitution and insertion variants of kmer at distance n_mismatch
      Look up each variant in index at distance n_mismatch
      Collect all matching reference positions → mapping_dict[n_mismatch]
    END FOR
    FOR n_mismatch = 0 TO max_n_mismatch DO
      IF mapping_dict[n_mismatch] is non-empty THEN
        Record positions and n_mismatch in mapping table for kmer
```

```
        BREAK (use best/lowest-mismatch match only)
      END IF
    END FOR
    Copy group and exclusivity labels from enriched table into mapping table
    IF reference gene annotations are defined THEN
      FOR each mapped position DO
        Find all genes whose annotated range covers that position
      END FOR
      Record union of gene names in mapping table for kmer
    END IF
END FOR
```

**STEP 3 — Compute per-position coverage**

```
FOR each kmer in enriched k-mers table DO
  FOR each start_position mapped to kmer DO
    Increment coverage counter for positions [start_position, start_position + k) in the column corresponding to kmer's group
  END FOR
END FOR
```

**STEP 4 — Save results**

```
Write mapping table to output/mapping_{k}mers_max_{max_n_mismatch}_mismatches.tsv
Write coverage table to output/coverage_{k}mers_max_{max_n_mismatch}_mismatches.tsv
```

**Construction of a pentapartite graph.** We hypothesized that the disjoint union of the "pentapartite graph"[95] can represent spatial interactions of isolates or enriched *k*-mers across different groups of HIV-1 subtypes. The "pentapartite graph"[95] is defined by the Equation below.

$$G = (V_1 \cup V_2 \cup V_3 \cup V_4 \cup V_5, E) \tag{7}$$

$$\text{where } E \subseteq \bigcup_{i=1}^{5} \{\{u, v\} : u, v \in V_i, u \neq v\} \tag{8}$$

$$\text{subject to: } V_i \cap V_j = \oslash \ \forall i \neq j \tag{9}$$

$$\text{and: } \{u, v\} \notin E \text{ if } u \in V_i \text{ and } v \in V_j \text{ for any } i \neq j \tag{10}$$

$V_i$ and $V_j$ are finite vertex sets, representing the union of the five groups of HIV-1 subtypes. Every vertex in the graph belongs to exactly one of the five groups. In this work, $V_i$ and $V_j$ represent either isolates (**Supplementary Fig. S4a-S4d**) or enriched *k*-mers (**Supplementary Fig. S4f-S4i**). $E$ is a finite set of edges, representing the sum of either "isolate *k*-mer count" (**Supplementary Fig. S4a-S4d**) or "subtype *k*-mer count" (**Supplementary Fig. S4f-S4i**) between two adjacent vertices in the same group of HIV-1 subtypes. The edges only connect vertices assigned to the same group $V_i$ or $V_j$, and no loops are allowed. The summation of each edge value was normalized by the sum of the total edge values in the same group of the HIV-1 subtype. In this work, undirected graphs are generated.

The matrices from the calculation of "isolate *k*-mer count" or "subtype *k*-mer count" were applied to construct a pentapartite graph. In each group of HIV-1 subtypes, we first indexed a matrix, followed by a random sampling of 100 isolates or enriched *k*-mers for the construction of a network in order to reduce computing cost. Here, it is important to stress that, given that PORT-EK-v2 retrieves enriched *k*-mers rather than unique *k*-mers present in each group of HIV-1 subtypes, the probability of biased distributions of enriched *k*-mers across all groups of HIV-1 subtypes resulting from presampling of a small size of the input dataset should be low. Although we cannot completely ignore a bias that can be introduced by missing edges between vertices in different groups of HIV-1 subtypes, our strategy ensures to focus on edges with statistical significance. We repeated this step 10 times (**Fig. 4a**).

We further allowed a maximum pairing of each vertex in every presampled small input dataset to generate a pairwise correlation matrix, in which the weighted edge was calculated based on the sum of either "isolate *k*-mer count" or "subtype *k*-mer count" between two adjacent vertices. Loops generated from the same vertices were excluded in this work. The weighted edge is defined in equation (4) in **Methods**. The pairwise distance between two adjacent vertices, defined in the following Equation, can then be computed.

$$D_{i,i'} = 1 - w_{i,i'} / \sum_{i=1}^{i} W \qquad (11)$$

, where $D_{i,i'}$ is the pairwise distance between isolate $i$ and isolate $i'$, $w_{i,i'}$ is the edge weight between isolate $i$ and isolate $i'$, and $W$ is the total sum of all values of edge weights computed in the same group of the HIV-1 subtype.

We utilized the function graph_from_data_frame() from the R package "igraph"[85] (https://igraph.org) with the following arguments: d for the edge list, vertices for the node list, and directed = FALSE to account for undirected edges and visualize in a network. Of note, we generated separate node lists for each input matrix. The computing code is written in R and available at GitHub (https://github.com/Quantitative-Virology-Research-Group/PORT-EK-version-2/tree/main/Analysis/Supplementary.figures).

**Calculation of assortativity coefficient.** Degree assortativity coefficients were computed using the function assortativity_degree() in the R package "igraph"[85] (https://igraph.org). The coefficient was computed based on 10 networks constructed by presampling 100 isolates (see the section "Construction of a pentapartite graph") for the calculation of statistical significance. Statistical tests were performed with R with default options.

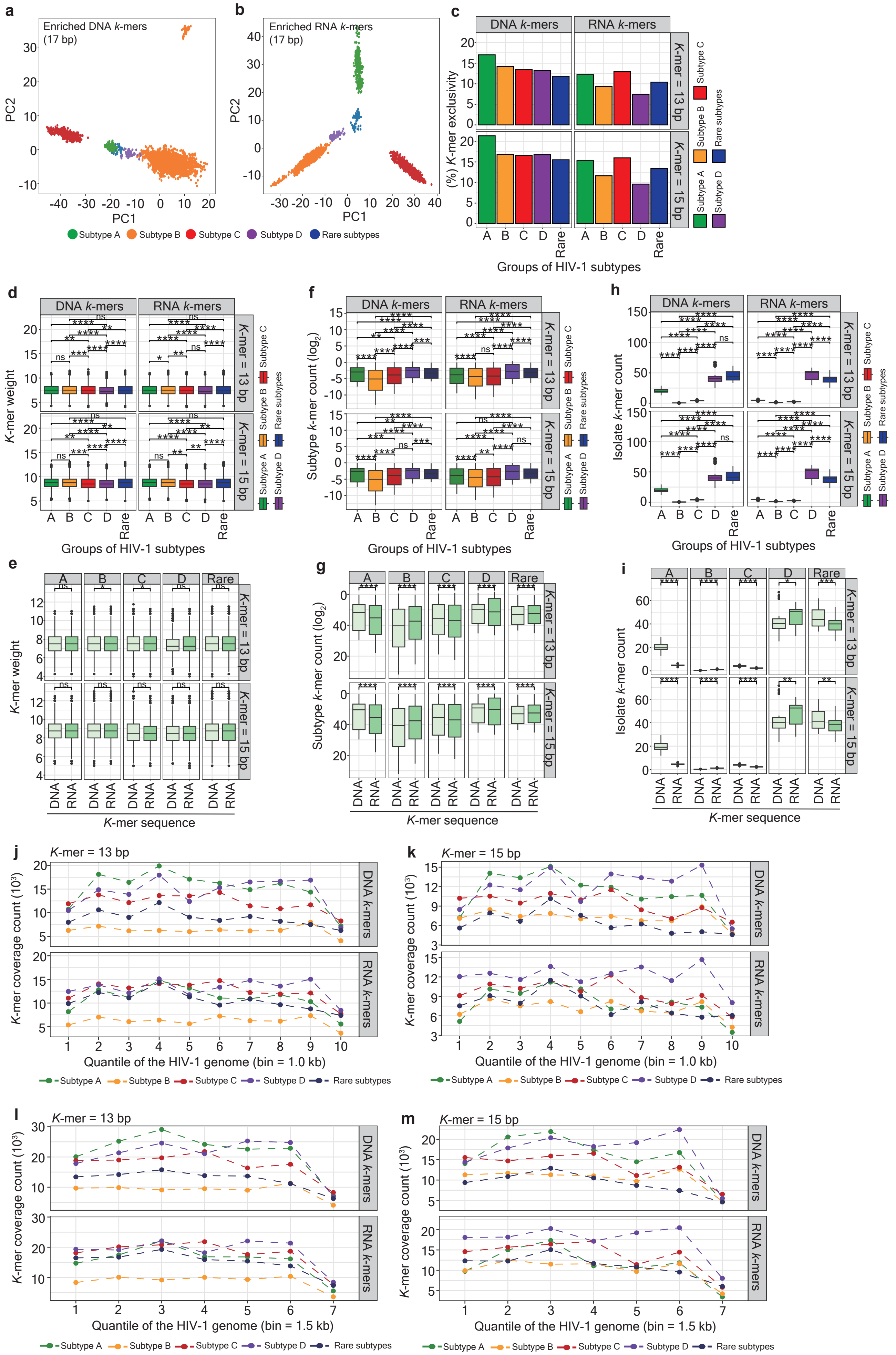
a
Enriched DNA k-mers
(17 bp)
PC2
PC1
b
Enriched RNA k-mers
(17 bp)
Subtype A
Subtype B
Subtype C
Subtype D
Rare subtypes
c
DNA k-mers
RNA k-mers
K-mer = 13 bp
K-mer = 15 bp
(%) K-mer exclusivity
Groups of HIV-1 subtypes
d
K-mer weight
e
K-mer sequence
f
Subtype k-mer count (log2)
g
h
Isolate k-mer count
i
j
K-mer = 13 bp
K-mer coverage count (10³)
Quantile of the HIV-1 genome (bin = 1.0 kb)
k
K-mer = 15 bp
l
m
Quantile of the HIV-1 genome (bin = 1.5 kb)

**Supplementary Fig. S1. Characterization of sequence patterns deciphered by enriched DNA and RNA *k*-mers.** (**a**, **d**) Two dimensional principal component analysis (PCA) revealing the discrepancy of enriched DNA (**a**) and RNA *k*-mers (**b**) in a length of 17 bp. Dots marked green, orange, red, purple, and dark blue represent HIV-1 subtype A, B, C, D, and the rare subtypes, respectively. (**c**) Bar charts representing the percentage of enriched *k*-mers, which are exclusively present in respective groups of HIV-1 subtypes. Boxes marked green, orange, red, purple, and dark blue represent HIV-1 subtype A, B, C, D, and the rare subtypes, respectively. (**d**, **f**, **h**) Box plots representing the percentage of "*k*-mer weight" (**d**), "subtype *k*-mer count" (**f**) displayed on a logarithmic scale, and "isolate *k*-mer count" (**h**) across different groups of HIV-1 subtypes. Facets at the x-axis separate enriched DNA and RNA *k*-mers; facets at the y-axis separate enriched *k*-mers in length of 13 bp or 15 bp. Boxes marked green, orange, red, purple, and dark blue represent HIV-1 subtype A, B, C, D, and rare subtypes, respectively. Significance levels are denoted as follows: ns for no significance, **p* 0.05, ***p* 0.01, ****p* 0.001, *****p* 0.0001. (**e**, **g**, **i**) Box plots representing the percentage of "*k*-mer weight" (**e**), "subtype *k*-mer count" (**g**) displayed on a logarithmic scale, and "isolate *k*-mer count" (**i**) between enriched DNA and RNA *k*-mers. Facets at the x-axis separate *k*-mers enriched from different groups of HIV-1 subtypes; facets at the y-axis separate enriched *k*-mers in a length of 13 bp or 15 bp. Significance levels are denoted as follows: ns for no significance, **p* 0.05, ***p* 0.01, ****p* 0.001, *****p* 0.0001. (**j**-**m**) Quantile plots representing the coverage of enriched *k*-mers in a length of 13 (**j** and **l**) or 15 bp (**k** and **m**) throughout the complete HIV-1 genome at the quantile of 1.0 kb (**j** and **k**) and 1.5 kb (**l** and **m**). Facets on the y-axis separate enriched DNA and RNA *k*-mers. Dots and dashed lines marked green, orange, red, purple, and dark blue represent HIV-1 subtype A, B, C, D, and the rare subtypes, respectively.

**a**

| | | Nucleotide | | | |
|---|---|---|---|---|---|
| | | A | T | C | G |
| Ordinal encoding condition | I | 0.25 | 0.50 | 0.75 | 1.00 |
| | II | 1.00 | 0.25 | 0.50 | 0.75 |
| | III | 0.75 | 1.00 | 0.25 | 0.50 |
| | IV | 0.50 | 0.75 | 1.00 | 0.25 |

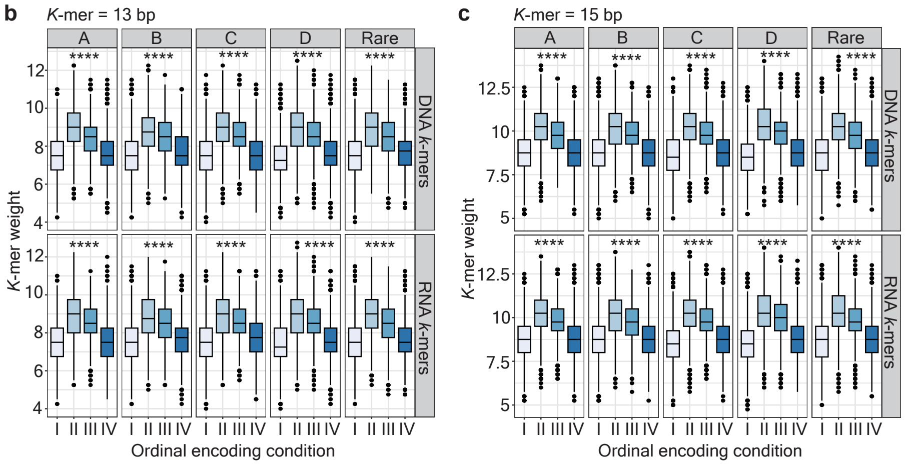


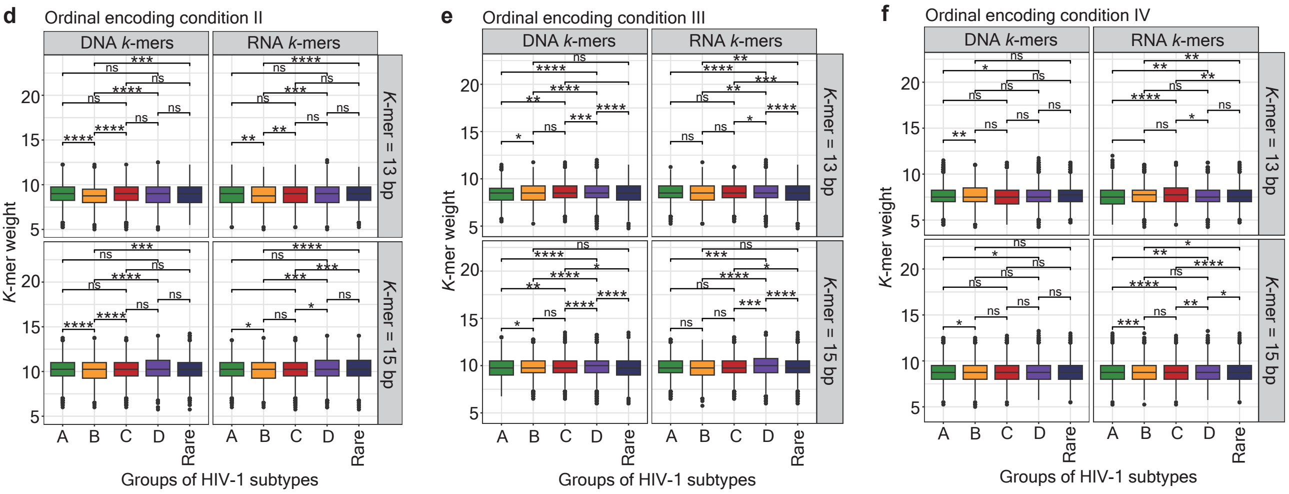


**Supplementary Fig. S2. Alteration of ordinal encoding towards nucleotides.** (**a**) Table representation of baseline encoding of four nucleotides in four ordinal encoding conditions. (**b**, **c**) Box plots representing a bias of "*k*-mer weight" computed from four different ordinal encoding conditions in a *k*-mer length of 13 bp (**b**) and 15 bp (**c**). Facets at the x-axis separate *k*-mers enriched from different groups of HIV-1 subtypes; facets at the y-axis separate enriched DNA and RNA *k*-mers. Significance levels are denoted as follows: ns for no significance, ****$p$ 0.0001. (**d**, **e**, **f**) Box plots representing "*k*-mer weight" computed based on the ordinal encoding condition II (**d**), III (**e**), and IV (**f**) across different groups of HIV-1 subtypes. Facets at the x-axis separate enriched DNA and RNA *k*-mers; facets at the y-axis separate enriched *k*-mers in length of 13 bp or 15 bp. Boxes marked green, orange, red, purple, and dark blue represent HIV-1 subtype A, B, C, D, and rare subtypes, respectively. Significance levels are denoted as follows: ns for no significance, *$p$ 0.05, **$p$ 0.01, ***$p$ 0.001, ****$p$ 0.0001. "*K*-mer weight" computed based on the ordinal encoding condition I is displayed in **Fig. S1d**.

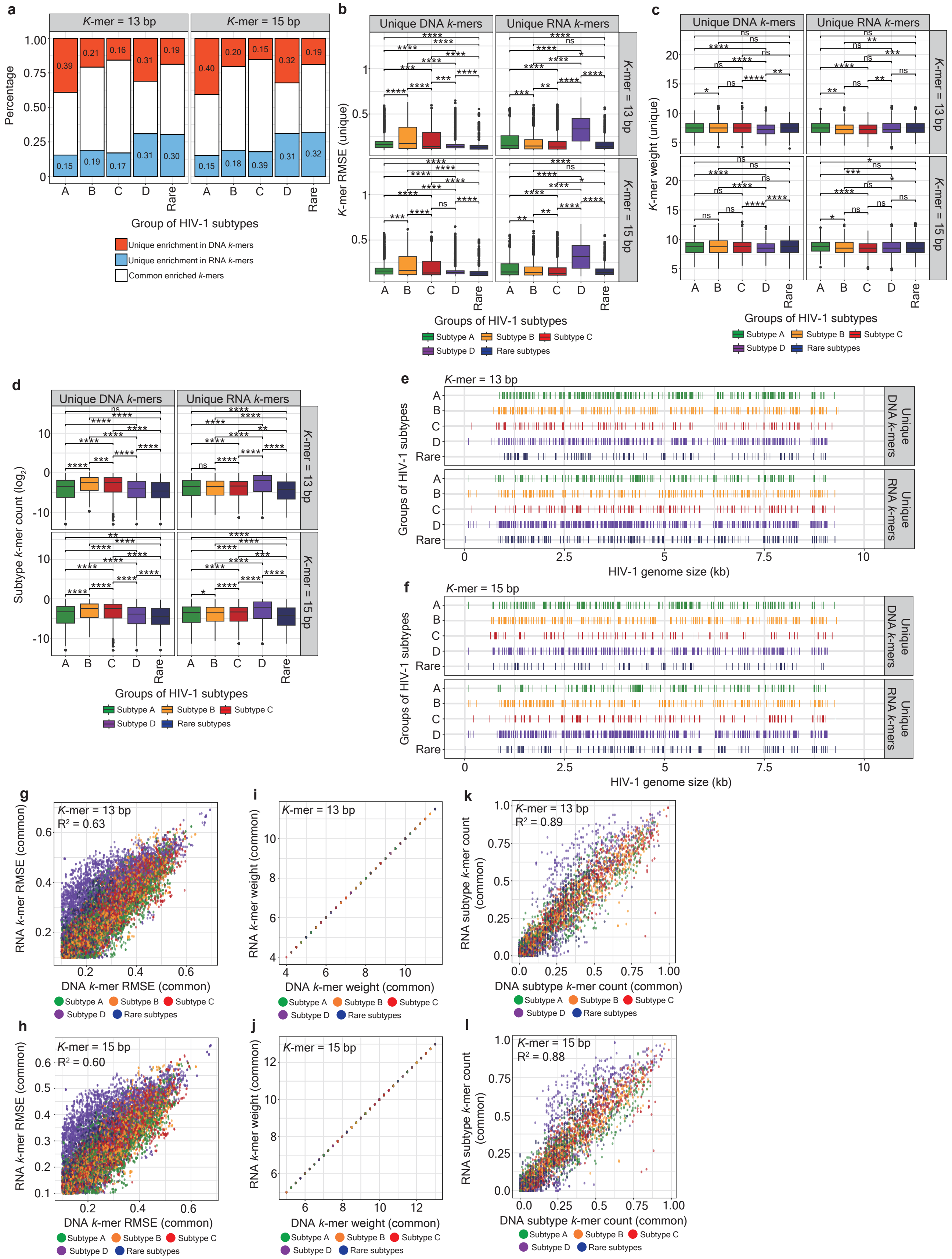

**Supplementary Fig. S3. Characterization of unique and common enriched DNA and RNA *k*-mers. (a)** Stacked bar charts representing the percentage of unique and common enriched DNA and RNA *k*-mers in a length of 13 bp (the left-hand side panel) and 15 bp (the right-hand side panel) across different groups of HIV-1 subtypes. Boxes marked red-orange, sky blue, and white represent unique enriched DNA and RNA *k*-mers and common enriched *k*-mers, respectively. **(b, c, d)** Box plots representing the percentage of "*k*-mer

RMSE" (**b**), "*k*-mer weight" (**c**), and "subtype *k*-mer count" (**d**) are displayed on a logarithmic scale across different groups of HIV-1 subtypes. Facets at the x-axis separate unique enriched DNA and RNA *k*-mers; facets at the y-axis separate enriched *k*-mers in length of 13 bp or 15 bp. Boxes marked green, orange, red, purple, and dark blue represent HIV-1 subtype A, B, C, D, and the rare subtypes, respectively. Significance levels are denoted as follows: ns for no significance, **p* 0.05, ***p* 0.01, ****p* 0.001, *****p* 0.0001. (**e**, **f**) Line plots visualizing the genomic loci, which present overlaps with unique enriched *k*-mers in a length of 13 bp (**e**) and 15 bp (**f**) across different groups of HIV-1 subtypes. Lines marked in green, orange, red, purple, and dark blue represent HIV-1 subtype A, B, C, D, and the rare subtypes, respectively. (**g-l**) Scatter plots representing the percentage of "*k*-mer RMSE" (**g** and **h**), "*k*-mer weight" (**i** and **j**), and "subtype *k*-mer count" (**k** and **l**) in a length of 13 bp (**g**, **i**, **k**) and 15 bp (**h**, **j**, **l**) across different groups of HIV-1 subtypes. Dots marked in green, orange, red, purple, and dark blue represent HIV-1 subtype A, B, C, D, and the rare subtypes, respectively. Of note, one tenth of the all common enriched *k*-mers were randomly retrieved to plot panels **g**, **h**, **k**, and **l**.

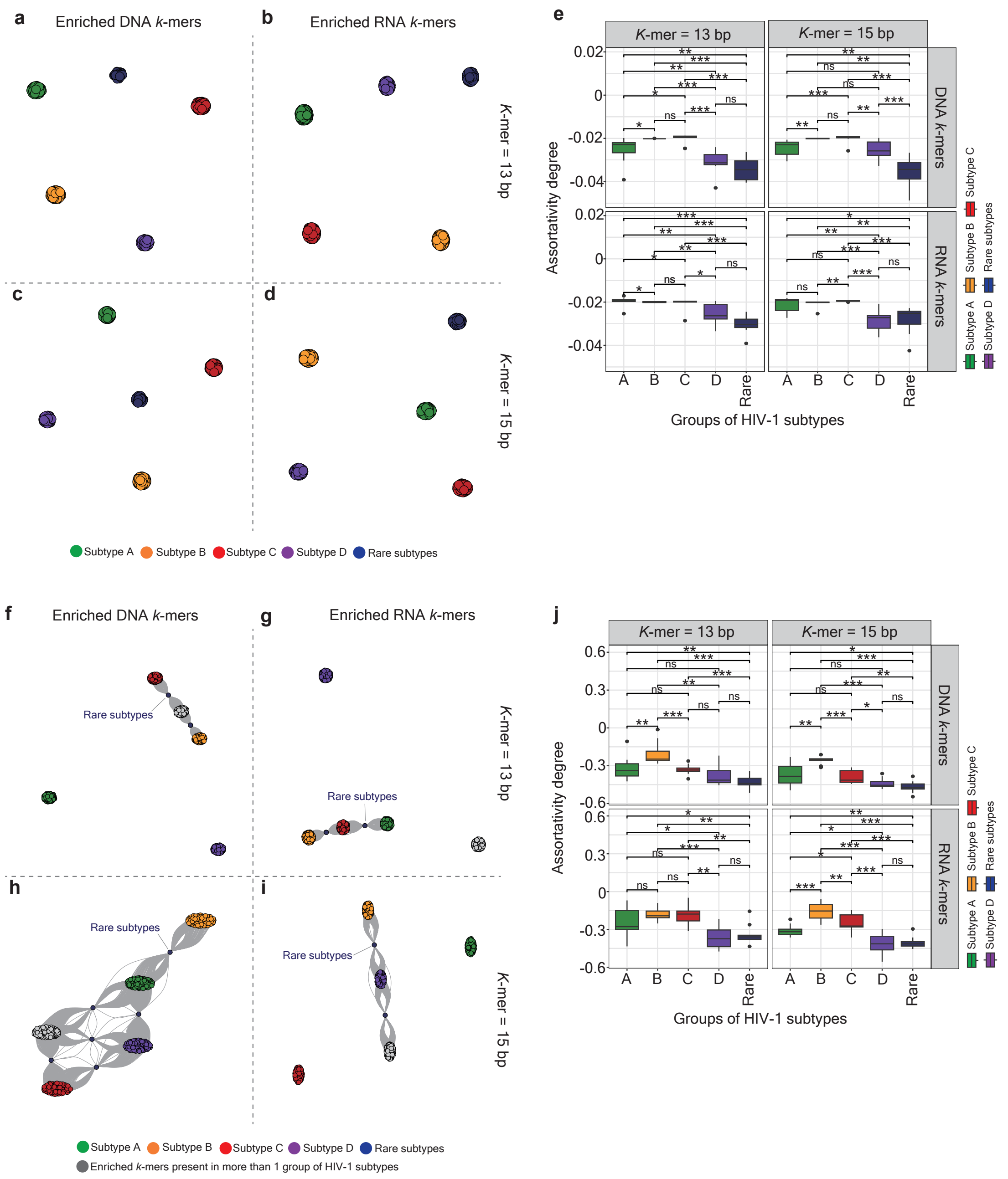


**Supplementary Fig. S4. Characterization of topological networks across different groups of HIV-1 subtypes.** (**a-d**) The pentapartite graphs illustrate the interactions between HIV-1 isolates across five different groups of HIV-1 subtypes. Networks were constructed by "isolate *k*-mer count" computed from enriched DNA (**a**, **c**) and RNA (**b**, **d**) *k*-mers in a length of 13 bp (**a**, **b**) and 15 bp (**c**, **d**), whereby each vertex represents an individual HIV-1 isolate. Vertices marked green, orange, red, purple, and dark blue represent HIV-1 subtype A, B, C, D, and the rare subtypes, respectively. (**e**) Box plots representing the degree assortativity coefficient of pentapartite graphs of networks constructed based on "isolate *k*-mer count" across different groups of HIV-1 subtypes. Facets at the x-axis separate *k*-mers in a length of 13 bp or 15 bp; facets at the y-axis separate enriched DNA and RNA *k*-mers. Boxes marked green, orange, red, purple, and dark blue represent HIV-1 subtype A, B, C, D, and the rare subtypes, respectively. Significance levels are denoted as follows: ns for no significance, *$p$ 0.05, **$p$ 0.01, ***$p$ 0.001. (**f-i**) The pentapartite graphs illustrate the interactions between two adjacent enriched *k*-mers across different groups of HIV-1 subtypes. Networks were constructed by "subtype *k*-mer count" computed from enriched DNA (**f**, **h**) and RNA (**g**, **i**) *k*-mers in a length of 13 bp (**f**, **g**) and 15 bp (**h**, **i**), whereby each vertex represents an individual enriched *k*-mer. Vertices marked green, orange, red, purple, and dark blue represent HIV-1 subtypes A, B, C, D, and rare subtypes, respectively; vertices marked grey represent enriched *k*-mers present in more than one group. (**j**) Box plots representing the degree assortativity coefficient of pentapartite graphs of networks constructed based on "subtype *k*-mer count" across different groups of HIV-1 subtypes. Facets at the x-axis separate *k*-mers in a length of 13 bp or 15 bp; facets at the y-axis separate enriched DNA and RNA *k*-mers. Boxes marked green, orange, red, purple, and dark blue represent HIV-1 subtype A, B, C, D, and rare subtypes, respectively. Significance levels are denoted as follows: ns for no significance, *$p$ 0.05, **$p$ 0.01, ***$p$ 0.001.

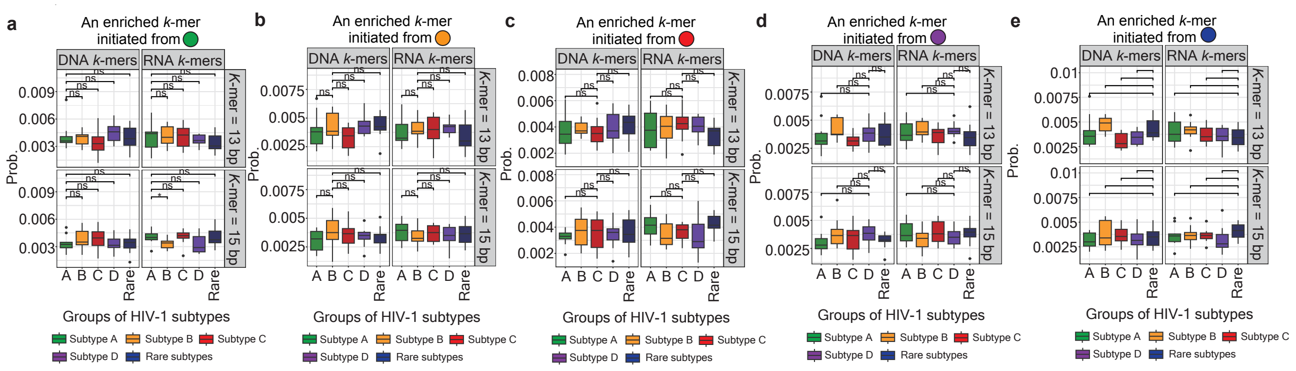


**Supplementary Fig. S5. The performance of MCMC modeling throughout networks constructed using mislabeled pairwise correlation matrices.** (**a**-**e**) Box plots representing the probability that vertices represent enriched *k*-mers to mislabled groups of HIV-1 subtypes within a path of each scenario, whereby a random walk initiates a vertex assigned to HIV-1 subtype A (**a**), B (**b**), C (**c**), D (**d**), and the rare subtypes (**e**). Networks were constructed by "subtype *k*-mer count", whereby each vertex represents an individual enriched *k*-mer. Facets at the x-axis separate enriched DNA and RNA *k*-mers; facets at the y-axis separate a *k*-mer length between 13 bp and 15 bp. Boxes marked in green, orange, red, purple, and dark blue represent HIV-1 subtype A, B, C, D, and the rare subtypes, respectively. Significance levels are denoted as follows: ns for no significance.

**Supplementary Table S1. Systematic comparison of parameters used between PORT-EK and PORT-EK-v2.**

| **Key parameters** | **Description** | **PORT-EK**[47] | **PORT-EK-v2** |
|---|---|---|---|
| *k* | *k*-mer length | Applied | Applied |
| *c* | Rarity filter threshold, range 0-1 | Applied | Set automatically based on *max_mem* instead |
| *max_mem* | Maximum memory size of *k*-mer count matrix in GB. | Not applied | Applied |
| *min_freq* | Calculation of minimum *k*-mer frequency (used only when count matrix size would surpass *max_mem*) | Not applied | Applied |
| *m* | Rare *k*-mers allowed mismatches, positive integer < *k* | Applied | Not applied |
| $min_{RMSE}$ | RMSE enrichment threshold, range 0-1 | Applied | Not applied |
| $m_{map}$ / *d* | Maximum mismatches allowed while mapping *k*-mers to reference, positive integer < *k* | Applied (one mapping location per *k*-mer). | Applied, name changed to *d* as it also covers indels. (multiple mapping locations possible) |
| $l_{map}$ | Maximum offset allowed while mapping *k*-mers to reference, positive integer | Applied | Not applied |

**Supplementary Table S10. AUC values recorded from logistic regression models. Related to Fig. 3a.**

| **Features** | ***K*-mer = 13 bp** | ***K*-mer = 15 bp** |
|---|---|---|
| Isolate *k*-mer count | 0.890 | 0.889 |
| Subtype *k*-mer count | 0.537 | 0.534 |
| *K*-mer average count | 0.576 | 0.577 |
| *K*-mer RMSE | 0.577 | 0.580 |
| *K*-mer weight | 0.523 | 0.521 |

**Supplementary Table S11. Probability recorded from multinomial logistic regression models. Related to Fig. 3b.**

| Groups of HIV-1 subtypes | | | | | | | | | | |
|---|---|---|---|---|---|---|---|---|---|---|
| | A | | B | | C | | D | | Rare | |
| ***K*-mer length (bp)** | | | | | | | | | | |
| | 13 | 15 | 13 | 15 | 13 | 15 | 13 | 15 | 13 | 15 |
| **DNA *k*-mers** | | | | | | | | | | |
| **Features** | | | | | | | | | | |
| Isolate *k*-mer count | 0.018 | 0.018 | 0.877 | 0.874 | 0.089 | 0.092 | 0.008 | 0.008 | 0 | 0 |
| Subtype *k*-mer count | 0.200 | 0.200 | 0.199 | 0.199 | 0.119 | 0.200 | 0.199 | 0.199 | 0 | 0 |
| *K*-mer average count | 0.249 | 0.249 | 0.152 | 0.153 | 0.206 | 0.201 | 0.248 | 0.256 | 0.143 | 0.139 |
| *K*-mer RMSE | 0.248 | 0.249 | 0.151 | 0.153 | 0.207 | 0.200 | 0.249 | 0.258 | 0.144 | 0.139 |
| *K*-mer weight | 0.241 | 0.243 | 0.151 | 0.153 | 0.205 | 0.202 | 0.249 | 0.256 | 0.144 | 0.138 |
| **RNA *k*-mers** | | | | | | | | | | |
| Isolate *k*-mer count | 0.141 | 0.141 | 0.512 | 0.512 | 0.310 | 0.311 | 0.020 | 0.019 | 0 | 0 |
| Subtype *k*-mer count | 0.199 | 0.199 | 0.200 | 0.199 | 0.199 | 0.200 | 0.200 | 0.199 | 0 | 0 |
| *K*-mer average count | 0.188 | 0.183 | 0.155 | 0.158 | 0.219 | 0.216 | 0.262 | 0.267 | 0.175 | 0.174 |
| *K*-mer RMSE | 0.188 | 0.183 | 0.153 | 0.158 | 0.219 | 0.217 | 0.260 | 0.266 | 0.175 | 0.174 |
| *K*-mer weight | 0.188 | 0.183 | 0.154 | 0.158 | 0.219 | 0.217 | 0.261 | 0.266 | 0.175 | 0.174 |

**Supplementary Table S12. Accuracy recorded from simple neural network-based models. Related to Fig. 3c.**

| | **DNA *k*-mers** | | **RNA *k*-mers** | |
|---|---|---|---|---|
| ***K*-mer length (bp)** | | | | |
| | 13 | 15 | 13 | 15 |
| **Features** | | | | |
| Isolate *k*-mer count | 95.419 | 95.731 | 76.310 | 78.105 |
| Subtype *k*-mer count | 0 | 0 | 0 | 0 |
| *K*-mer average count | 26.503 | 24.996 | 31.306 | 33.762 |
| *K*-mer RMSE | 26.386 | 27.363 | 31.306 | 32.383 |
| *K*-mer weight | 25.232 | 25.958 | 24.952 | 27.309 |
